# FLASHIda: Intelligent data acquisition for top-down proteomics that doubles proteoform level identification count

**DOI:** 10.1101/2021.11.11.468203

**Authors:** Kyowon Jeong, Maša Babović, Vladimir Gorshkov, Jihyung Kim, Ole N. Jensen, Oliver Kohlbacher

## Abstract

Top-down proteomics (TDP) has gained a lot of interest in biomedical application for detailed analysis and structural characterization of proteoforms. Data-dependent acquisition (DDA) of intact proteins is non-trivial due to the diversity and complex signal of proteoforms. Dedicated acquisition methods thus have the potential to greatly improve TDP. We present FLASHIda, an intelligent online data acquisition algorithm for TDP that ensures the real-time selection of high-quality precursors of diverse proteoforms. FLASHIda combines fast charge deconvolution algorithms and machine learning-based quality assessment for optimal precursor selection. In analysis in *E. coli* lysates, FLASHIda increased the number of unique proteoform level identifications from 800 to 1,500, or generated a near-identical number of identifications in ⅓ of instrument time when compared to standard DDA mode. Furthermore, FLASHIda enabled sensitive mapping of post translational modifications and detection of chemical adducts. As an extension module to the instrument, FLASHIda can be readily adopted for TDP studies of complex samples to enhance proteoform identification rates.

## Main

Top-down proteomics (TDP) enables comprehensive and in-depth analysis of intact proteoforms (i.e., protein species arising from the same gene product via splice variants, genomic variation, post-translational modifications, degradation, etc.^1, 2^). Proteoforms have high phenotypic heterogeneity in different biological systems, and proteoform level information can provide important insights into biochemical function or disease phenotypes^3–7^. TDP as the method of the choice to study proteoforms has thus become important for many biomedical applications^8, 9^.

In recent years, significant improvements have been made to sample processing, separation, fragmentation, and bioinformatics methods for TDP^10–15^. As a result, proteoforms that have been difficult to analyze by TDP (e.g., proteoforms of large masses and of membrane proteins) have become easier to detect and characterize^7, 8^. In large-scale studies of complex samples such as microbial or human cell lysates, the number of proteoform identification has been increased up to 4,000-6,000 (corresponding to 500-1,000 proteins)^16, 17^. In single-shot TDP experiments, about 800 proteoforms could be identified in *E. coli* lysate^18^ and about 1,800 proteoforms in a human brain sample^19^. The broad coverage in these experiments can be attributed to technical improvements like capillary zone electrophoresis (CZE) and high-field asymmetric waveform ion mobility spectrometry (FAIMS)^19–22^ as well as the state-of-the-art bioinformatics analysis tools like TopPIC^14^ and ProSight PC (Thermo Fisher).

At the same time, fragmentation strategies implemented in data-dependent acquisition (DDA) by current instrument software have been optimized to support bottom-up proteomics (BUP) rather than for TDP. These strategies usually select *N* most intense peaks (typically, *N* is in a range of 10 - 20) for fragmentation acquisition, often excluding previously selected ones over a short retention time (RT) period (Top-*N* acquisition with a dynamic exclusion list)^23^. While this scheme effectively captures diverse peptide ions of high quality in BUP studies^24^, these selection criteria are inadequate for optimal proteoform ion selection. In contrast to peptide ions in BUP, a single proteoform generates many peaks due to its large mass and high charges. Top-*N* acquisition thus often results in the selection of multiple peaks from a single abundant proteoform rather than from multiple distinct proteoforms, which in turn results in poor proteoform coverage. In addition, the exclusion list preventing the selection of the same m/z (mass-to-charge ratio) region does not guarantee the exclusion of the same proteoform (mass).

Most large-scale TDP studies, however, use DDA acquisition with specifically tuned parameters, for instance, relatively low *N* values (ranging from 3 to 5) for Top-*N* acquisition and widened isolation window width up to 5-15 Th^5, 18^. Post hoc analysis of the selected precursor ions shows that the selection of the proteoforms is far from ideal. Smarter acquisitions, sometimes termed ‘intelligent data acquisition’ (IDAs), have thus been discussed in the literature (e.g., Durbin et al.^25^ and Lu et al.^11^) using real-time mass deconvolution (i.e., real-time determination of intact proteoform masses) to disentangle complex signal structure of proteoforms. Autopilot^25^ aimed to divertize the selected proteoform masses, and MetaDrive^11^ focused on boosting the quality of MS2 spectra by merging multiple precursors from the same proteoform. Both approaches clearly demonstrated the potential of IDA for TDP studies and, at the same time, were limited by the need for faster mass deconvolution methods showing only 3-5% improvement upon standard DDA acquisition in proteoform identification count.

Here, we present FLASHIda, a novel machine learning-based IDA algorithm designed to maximize proteoform coverage in TDP. FLASHIda interfaces with tribrid Thermo Scientific mass spectrometers through the instrument API (iAPI) allowing for real-time access to MS data. By combining our recently developed real-time spectrum deconvolution algorithms^26^ and a machine learning technique assessing the quality of the precursor isotopomer envelopes, FLASHIda enables non-redundant selection of precursor ions of very high quality during the LC-MS run and thus boosts the proteoform coverage.

## Results

### FLASHIda overview

Fig. 1a illustrates the MS duty cycle control employed by FLASHIda. FLASHIda processes each MS full scan within a few milliseconds (about 20 ms on average) and optimizes the acquisition of the next cycle to maximize isoform diversity in acquisition (see Methods for the algorithm). Fig. 1b illustrates the key steps of FLASHIda. FLASHIda uses Thermo iAPI to access MS full scan in real-time. FLASHIda takes two steps to select high quality precursor isotopomer envelopes (simply precursors from here on) of diverse proteoforms. The first step is to transform the input m/z-intensity spectrum into a mass-quality spectrum, and the second is to select precursors in the transformed spectrum so that the number of uniquely identified proteoform masses is maximized.

**Figure 1.**
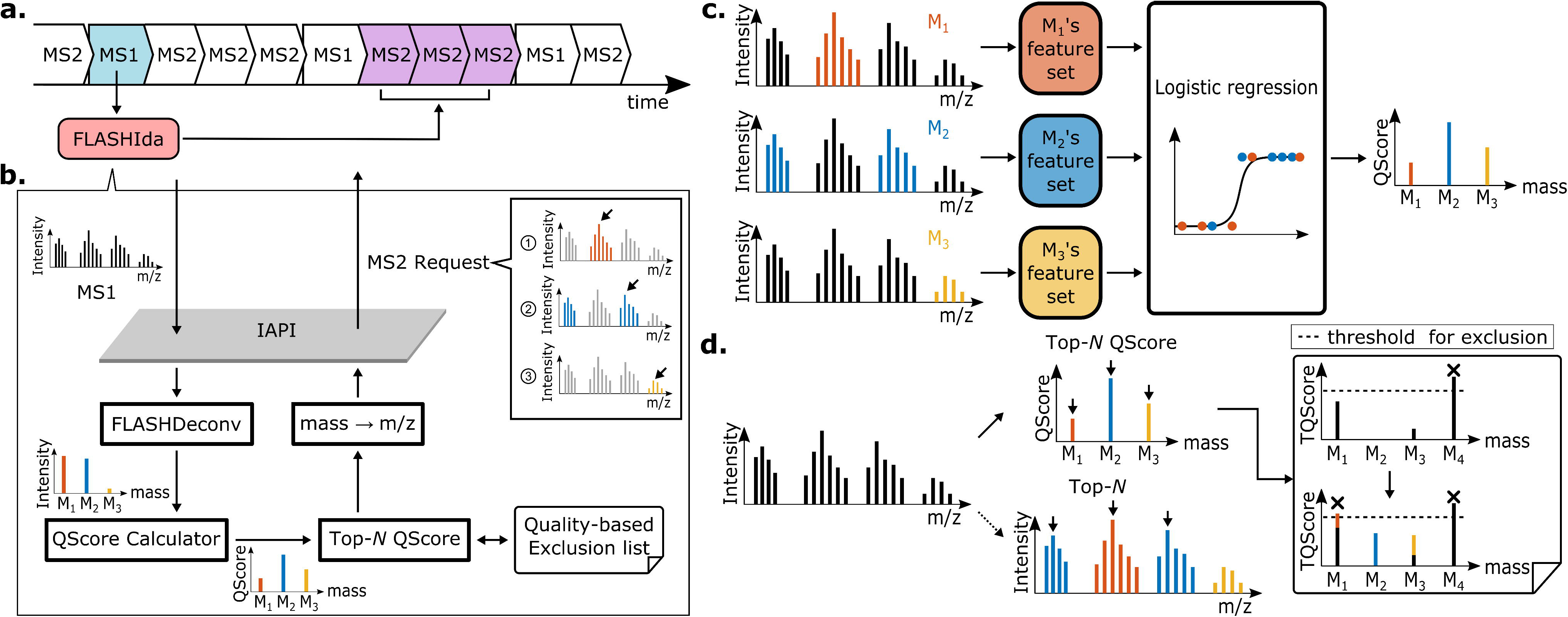
Overview of FLASHIda. **a.** MS duty cycle control employed by FLASHIda. FLASHIda processes each MS full scan within a few milliseconds (about 20 ms on average) and optimizes the acquisition of the next cycle to maximize isoform diversity in acquisition. **b.** Key steps of FLASHIda. FLASHIda uses Thermo iAPI to access MS full scan in real-time. FLASHIda takes two steps to select high quality precursor isotopomer envelopes of diverse proteoforms. The first step is to transform the input m/z-intensity spectrum into a mass-quality (QScore) spectrum, and the second is to select precursors in the transformed spectrum so that the number of unique proteoform level identifications (simply proteoform ID) is maximized (Top-*N* QScore precursor acquisition with a quality-based exclusion list). The selected precursor envelope mass ranges are converted back into m/z ranges (isolation windows) for MS2 acquisition through Thermo iAPI interface. **c.** QScore calculation via logistic regression. After the deconvolution, the quality of the resulting fragmentation is measured for each precursor using a logistic regression with six features (see Methods). The features are extracted from the peaks in the original spectrum corresponding to the precursor. The quality metric, or QScore, is the probability estimated by the regression that the resulting fragment spectrum to be successfully identified. The calculated QScores of masses are used instead of intensities of masses. **d.** Top-*N* QScore precursor selection with a quality-based exclusion list. With the transformed mass-QScore spectrum, FLASHIda attempts to select the high quality precursors while maximizing the identified proteoform diversity. The selection of high quality precursors is done by simply taking the top *N* masses with the highest QScore (Top-*N* QScore acquisition). To divertize the proteoform IDs, FLASHIda conservatively excludes masses that are highly likely to be already identified; if a mass has been acquired multiple times within a short RT duration, the probability that at least one of the acquired MS2 spectra from the mass is identified is calculated with its QScores (see Methods for detail). This probability, called TQScore, is immediately updated per mass upon each acquisition, and the masses of high TQScores (>0.9) are excluded from the acquisition for a short RT duration.

### Spectrum deconvolution and quality prediction

On receiving MS1 full scan, FLASHIda performs an instant mass deconvolution based on the fast decharging algorithm developed for FLASHDeconv^26^. Since a single (monoisotopic) mass is represented by multiple isotopomer envelopes of distinct charges, this transformation drastically reduces the signal complexity, already making the selection of distinct proteoform masses far simpler than in the original spectrum.

We then employ a machine learning model to predict the quality of the resulting fragmentation (Fig. 1c). Based on six relevant features (see below and Methods) extracted from the original mass spectrum, we have trained a logistic regression model to compute a quality metric (termed QScore), which predicts a probability for the resulting fragment spectrum to be successfully identified. Note that this does not require an actual identification, but merely estimates a likelihood for success based on the observation that higher ‘quality’ of the precursors affects the quality of the fragment spectrum and thus the identification likelihood. In preliminary studies, we found that the following features are easy to extract (speed is important for the online processing) and still contain sufficient information to assess the precursor quality: the shape of isotopomer envelopes (as compared to theoretical ones), signal-to-noise ratio (SNR) within the precursor envelope range, intensity distribution over different charges, and average mass errors of all peaks. The first two features (the isotopomer envelope shape and SNR) are defined both for peaks within precursor ion m/z range and for peaks within precursor ion mass range (of all possible charges). Detailed regression procedure and definition of the features are given in Methods. The calculated QScores of precursor masses can then be used to prioritize precursor selection in a manner that favors precursors with better odds of identification.

### Precursor prioritization

Once the quality scores of precursors have been computed, FLASHIda selects the best precursors from all available masses. To maximize the proteoform diversity, high-quality precursors from yet unfragmented proteoforms should be selected. The selection of high-quality precursors can be readily done based on the QScore of the precursors, simply by taking the top *N* masses with the highest QScore that have not yet been selected (Fig. 1d). In order to avoid re-fragmentation of the same proteoforms, we apply dynamic exclusion (inspired by the dynamic exclusion used in BUP) to maximize the diversity of the selected proteoforms.

We found that the number of high quality proteoform precursors at a given RT is often very low as compared with that of peptide precursors in BUP. Therefore, the use of the naive mass exclusion alone resulted in the selection of low quality masses, and in turn an overall drop of identification rate (see Supplementary Fig. 1). Instead, FLASHIda only excludes masses that are highly likely to be already identified. To this end it keeps a list of the triggered masses over a short RT duration along with their QScores. If a mass has been acquired multiple times, the probability that at least one of the acquired MS2 spectra from the mass is identified is calculated with its QScores (see Methods for details). Note that this probability, called TQScore (Total QScore) of the mass, is the probability that the proteoform of the mass is identified. After each acquisition, TQScores are immediately updated. If the TQScore of a mass exceeds 0.9, the mass is registered in the exclusion list for a short RT duration. Using this quality-based mass exclusion list, FLASHIda assures each triggered mass (of proteoform) is identified while still maximizing the diversity of proteoforms.

### IDA doubles the number of proteoform IDs compared to DDA

To benchmark FLASHIda, we generated four sets of nano-RPLC MS/MS single-runs from *E. coli* lysate: two sets using FLASHIda acquisition, and two using standard acquisition for intact proteins (Intact Protein Mode) on a Thermo Scientific Orbitrap Eclipse (see Methods for details). One set from each acquisition has a gradient time of 30 min and another 90 min. Each set was measured in technical triplicate, thus a total twelve measurements were obtained. The triplicates of 90 and 30 min FLASHIda runs are collectively denoted as FI90s and FI30s, respectively. Likewise, the standard runs are ST90s and ST30s. FI90s and FI30s are collectively called FI, and likewise ST. We note that the datasets used to train QScore are not part of this benchmark data.

In the benchmark tests, we distinguish proteoform level identifications, or simply proteoform IDs, and biological proteoforms since multiple proteoform IDs in the search results may represent simple chemical adducts of the same biological proteform.

For acquisition (both FLASHIda and standard), we used Top-4 acquisition and set precursor charge range between 4 and 50. For standard, dynamic exclusion with 30 s exclusion duration was used. For FLASHIda, mass range was set to from 400 to 50,000 Da, and the TQScore threshold was 0.9. In addition, FLASHIda discarded any precursors with QScore lower than 0.25 so only high quality precursors are triggered. Lastly, FLASHIda also discarded precursors of SNR within the precursor envelope m/z range (called precursor SNR) lower than 1.0 to minimize both precursor ion coelution and mass artifacts that result in excessive false positive proteoform IDs. Briefly, the MS2 spectra from coeluted precursor ion (also known as chimera spectra^27^) and of incorrect precursor masses often cause target-decoy based false discovery rate (FDR) control inaccuracies and even erroneously inflate the number of proteoform IDs (see Methods, Supplementary Fig. 2-3 and Supplementary Table. 1 for detail). The precursors of low precursor SNR often correspond to coeluted precursors or precursors of harmonic mass artifacts and thus often have incorrectly determined masses (see examples in Supplementary Fig. 4-8). Therefore, such precursors were avoided in data acquisition.

For data analysis, the twelve LC-MS/MS datasets were analyzed in the same way: raw files were converted into mzML files^28^ using ProteoWizard^29^ msconvert (version 3.0.20186-dd907d75; see Supplementary Fig. 9 for GUI screenshot), deconvoluted by FLASHDeconv (version 2.0 beta+), and identified by TopPIC (version 1.4.4). For FLASHDeconv, charge range was set to a range from 4 to 50 and mass range from 400 and 50,000 Da (identical with FLASHIda acquisition parameters). As for the acquisition, the precursor SNR threshold was set to 1.0 for data analysis; this threshold was already applied for FI datasets by FLASHIda but not for ST datasets. For TopPIC, one unknown modification was allowed for search, and the FDR threshold was set to 1% (both in spectrum and proteoform levels). We used the *E. coli* (strain: K12 MG1655 i) database in fasta format downloaded from SwissProt (downloaded 28.05.2020). The FLASHDeconv and TopPIC commands are shown in Methods. Fig. 2a shows the number of unique proteoform IDs, proteins, and spectra. FI90s resulted in nearly 1,500 proteoform IDs on average, almost doubling the number of proteoform IDs found in SI90s (792 on average). From FI30s, on average 764 proteoform IDs have been found, which is comparable to SI90s although FI30s used only a third of the machine time compared to SI90s. SI30s yielded the minimum proteoform ID count of 422 on average. In terms of protein count, FI90s reported 338 average protein count while SI90s reported 259. FI30s also reported 247 proteins on average, again demonstrating the high performance of FLASHIda in short gradient runs. ST30s reported about 181 proteins. The right panel of Fig. 2a compares the number of acquired, deconvolved, and identified precursors (or MS2 spectra). The numbers of identified MS2 spectra in FI datasets were almost twice as many as in ST, resulting in a rate of ~75% of fragment spectra being identifiable in FLASHIda runs, in contrast to ~35% using DDA. These results show that FLASHIda successfully selects high quality precursors from diverse proteoform IDs, significantly increasing the number of proteoform IDs (see also Supplementary Fig. 10-12 for more spectrum level analyses).

**Figure 2.**
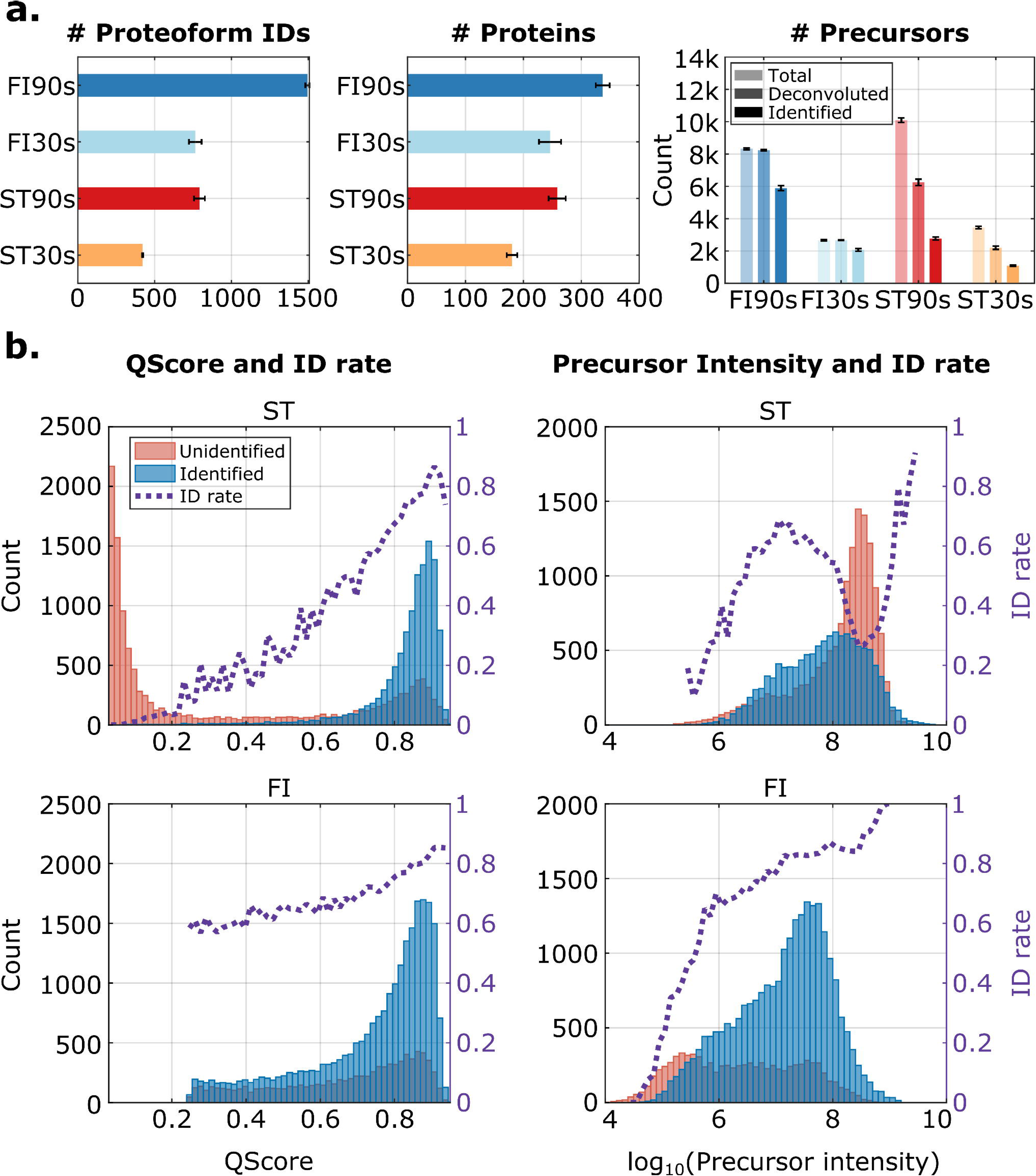
**a.** Identification results from FLASHIda acquisition as compared with the standard acquisition. Four sets of *E. coli* lysate single runs were prepared: 90 minute gradient (technical) triplicates with FLASHIda acquisition (FI90s), 30 minute FLASHIda (FI30s), 90 minute standard acquisition (ST90s), and 30 minute standard (ST30s). Top-4 acquisitions were used for both methods. Each dataset was deconvoluted by FLASHDeconv and identified by TopPIC at 1% spectrum and proteoform level FDR (see Methods). The left panel shows the unique proteoform ID count from each set, and the middle panel the protein count. The right panel shows the numbers of total triggered, deconvoluted, and identified precursors (or MS2 spectra) from each set. The error bars show the standard deviations. It is clearly shown that FLASHIda reports almost twice more unique proteoform IDs than the standard acquisition, or alternatively similar numbers as with standard acquisition on drastically shorter gradient runs. In terms of protein count, almost 30% increase was observed. The spectrum level analysis in the right panel shows that FLASHIda runs result in far better identification rates (~75%) than standard (~35%). **b.** The histograms for identified (blue) and unidentified (red) spectra over QScores (left panel) and precursor intensities (right) from the standard acquisition (ST, top panel) and FLASHIda (FI, bottom). For each plot, the identification rate (ID rate) in each bin is also shown as a purple dotted line. For FLASHIda runs, the distribution over QScore is truncated as precursors of QScore less than 0.25 were discarded. The top panel (ST) shows that QScore not only far better separates between identified and unidentified spectra than precursor intensity but also estimates the identification rate well in standard runs. In FLASHIda runs, QScore is underestimating the identification rate, which should be because QScore was used for the acquisition making the prediction biased. Comparing precursor intensity distributions also reveals that FLASHIda achieves almost an order of magnitude wider dynamic range than the standard acquisition.

Next, we assessed the prediction accuracy of the precursor quality score. Fig. 2b shows the histograms of identified and unidentified spectra with respect to QScore (left panel) and precursor intensity (right) from ST (top) and FI (bottom) datasets. The QScore distributions from ST datasets demonstrate that QScore separates the identified and unidentified ones far better than intensity. In addition, it is shown that QScore predicts the identification rate accurately. In FI datasets, the histogram is truncated and does not show low QScores, as precursors with values below 0.25 are discarded in IDA. Also the identification rate is underestimated, which should be because QScore was used for the acquisition rendering the prediction biased (QScore was trained on standard DDA acquired data). Comparing precursor intensity distributions (Fig. 2b, right panel) also reveals that FLASHIda achieves almost one order of magnitude higher dynamic range than standard DDA, suggesting that FLASHIda enables the selection of low-intensity, but high-quality precursors.

The advantage of FLASHIda acquisition is maximized when coupled with FLASHDeconv in the analysis because the precursor masses from both tools are consistent with each other. When FI and ST datasets were deconvolved by TopFD^14^ (for both MS1 and MS2) instead of FLASHDeconv, we first found that only 37.6% of distinct nominal proteoform ID masses from TopFD overlap with those from FLASHIda. Out of the non-overlapping proteoform IDs from TopFD, about 80% had precursor SNR lower than 1.0. After precursor SNR filtration (threshold of 1.0), the percentage of the overlapping nominal proteoform ID masses increased to 69.3% from 37.6%. Supplementary Fig. 13 and Supplementary Table 2-6 provide TopFD identification results, and Supplementary Fig. 14-18 provides the analyses on the TopFD proteoform IDs with low precursor SNRs. In terms of the identification count, only 5-10% more proteoform IDs were found in FI datasets than in ST datasets regardless of discarding proteoform IDs of low SNR precursors or not. When the precursor masses from FLASHIda were used instead of TopFD precursor masses (with MS2 deconvolution still done by TopFD), the boost was recovered to 50-65% (see Supplementary Table 7). Since MS2 deconvolution usually is dependent on precursor mass and charge values, the boost could be even higher if an interface between FLASHIda and TopFD is implemented.

### IDA improves reproducibility of proteoform level identification

Then we measured the proteoform and protein level reproducibility across all FI and ST datasets. Two proteoform IDs (from different datasets) are defined to match each other if they have the same amino acid sequence and masses within 1.2 Da tolerance. The overlap coefficients (the size of the intersection divided by the smaller of the size of two compared sets) between the datasets are given in Fig. 3a. Even with significantly larger numbers of proteoform IDs and proteins than ST datasets, FI datasets showed comparable overlap coefficients with ST datasets. For both datasets, proteoform level reproducibility (0.5-0.8) was rather lower than protein level (~0.8). As seen in Supplementary Fig. 19, proteoform reproducibility is more strongly correlated with precursor intensity than with quality (QScore). We thus presume that we are hitting the limits of stochasticity of detectability, which renders the low proteoform level reproducibility. Indeed, in many cases more than tens of proteoform IDs from a single gene (or protein) were identified with a high dynamic range. For example, 40 proteoform IDs were reported from DNA-binding protein H-NS (UniProtKB: P0ACF8) in a single FI90 dataset with four orders of magnitude dynamic range. Thus, for an overlapping protein, often some of its low abundant proteoform IDs were not detected across all datasets.

**Figure 3.**
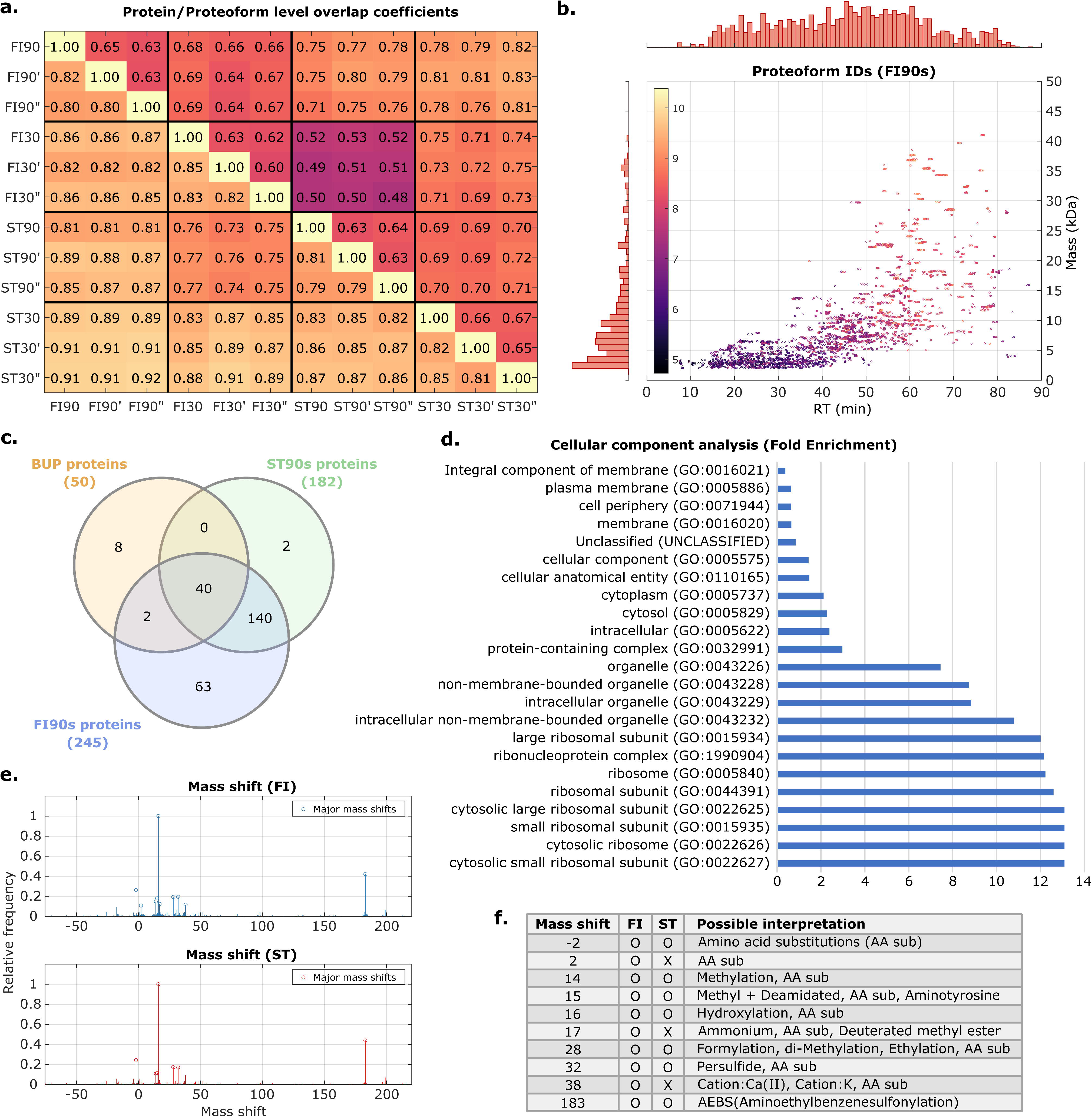
**a.** Protein (lower diagonal) and proteoform (upper) level overlap coefficients over all datasets. FI90, FI90’, and FI90” denote the triplicate datasets of FI90s, and the same notation applies for the other datasets. Two proteoform IDs from distinct datasets were defined to match each other when they have the same amino acid sequence and their mass difference is less than 1.2 Da. Even with significantly larger numbers of proteoform IDs and proteins than standard acquisition datasets, FLASHIda datasets showed comparable overlap coefficients with the standard datasets. **b.** Scatter plot showing the proteoform IDs from FI90s datasets over retention time (RT) and mass. In the top and left, marginal histograms over RT and mass are shown. Each proteoform ID is color coded by its intensity (measured by its feature area). Most proteoform IDs are present in the low mass region (<20 kDa), demonstrating that identifying large proteoforms via TDP is still difficult. The proteoform IDs are sparsely distributed over RTs from 60 to 90 minutes. The dynamic range of proteoform intensities is almost 6 orders of magnitude. **c.** Comparison between the identified proteins in FI90s and ST90s datasets with 50 high abundant *E. coli* proteins reported in the BUP study by Schmidt et al.^30^. From FI90s, 245 proteins jointly detected in all replicates were collected. Likewise, 182 proteins were collected from ST90s triplicates. Out of the 50 proteins from BUP, 42 and 40 were identified in FI90s and in ST90s datasets, respectively, showing high quantitative consistency although our results are from TDP. Six of the eight undetected proteins were heavy proteins (>35 kDa; see Supplementary Table 8). It is also shown that almost all the proteins (~99%) in ST90s datasets are also found in FI90s while 26% of FI90s proteins are exclusive. **d.** Cellular component gene ontology (GO) analysis for the proteins identified from FI90s triplicates, using the Gene Ontology Resource^31, 32^. Binomial tests at 5% false positive rates were used. The most abundant proteins are ribosomal proteins, as reported in Ishihama et al.^33^, but their proteoform level dynamic ranges varied for different proteins up to two orders of magnitude. Relatively rare membrane proteins were reported, indicating that analyzing intact membrane proteins via TDP is still a challenge. See Supplementary Table 9 for more GO analysis. **e.** Histograms for mass shifts in identified proteoform IDs from all FLASHIda runs (FI; top panel) and standard (ST; bottom). The mass shifts with relative frequency higher than 10% are considered as major ones and marked with circles. For both acquisition methods, the mass shifts were highly consistent with each other. **f.** Possible interpretations of major (circled) mass shifts found in **e**. The second and third columns specify if the mass shift is reported (as major ones) in FLASHIda runs (FI; the second column) and in standard (ST; the third). We referred to Unimod^46^ for possible interpretations of mass shifts in the last column.

### Analysis of proteoform and protein coverage

We then analyzed the proteoform IDs reported in the FI90s datasets. All 4,483 proteoform IDs are shown on the scatter plot on the RT-mass plane in Fig. 3b. The deconvoluted mass signal is shown to be rather sparse in particular in the later part of the gradient (60-80 min) where larger proteoforms elute. This would be because of low ionization efficiency and/or low signal (mainly consisting of isotopically unresolved peaks) quality of heavy proteoforms (>30 kDa). However, as compared to other *E. coli* TDP studies, far more heavy proteoform IDs were identified. For example, in a single FI90 dataset, 105 proteoform IDs with mass larger than 30 kDa have been identified (7% out of all proteoform IDs) while only 52 were identified (0.1% out of all) in a large-scale TDP analysis done in McCool et al.^17^ (2D LC separation, 43 CZE-MS/MS runs). The dynamic range of the proteoform intensities (measured by the feature area of each proteoform ID) is about five to six orders of magnitude.

Then, we compared the identified proteins in FI90s and ST90s datasets with published quantitative BUP data for *E. coli* MG1655 cells grown in LB medium from the study by Schmidt et al.^30^. From FI90s, 245 proteins jointly detected in triplicates were collected. Likewise, 182 proteins were collected from ST90s triplicates. Out of the 50 proteins reported to have the highest copy number per cell, 42 and 40 were identified in FI90s and in ST90s datasets, respectively (Fig. 3c). Six of the eight undetected proteins were heavy proteins (>35 kDa; see Supplementary Table 8). Fig. 3c also shows that almost all the proteins (~99%) in ST90s datasets are also found in FI90s while 26% of FI90s proteins are not found in ST90. We performed a gene ontology (GO) term analysis for the identified proteins, using the Gene Ontology Resource^31, 32^. Binomial tests at 5% false positive rates were used. The cellular component analysis (in Fig. 3d; also see Supplementary Table 9) shows that most abundant proteins are ribosomal proteins, as reported in Ishihama et al.^33^, another *E. coli* study done by BUP. On the proteoform level as well, almost 40% of identifications were from ribosomal proteins: 28% from 50S and 12% from 30S subunits. However, the number of proteoform IDs per ribosomal protein varied widely; for example, while a single unmodified proteoform ID of 50S ribosomal protein L15 (UniProtKB: P02413) was reported, 28 proteoform IDs of 50S ribosomal protein L31 type B (UniProtKB: P0A7N1) were identified in a single FI90 dataset, excluding six low-quality identifications (see Supplementary Table 10). The relative intensities of the ribosomal proteoform IDs from the same protein were also often two orders of magnitude apart (e.g., UniProtKB: P0A7N1, P0A7K2, and P0A7N9). We can thus confidently state that FLASHIda is able to provide rich information even for families of proteoforms including low-abundance species.

In addition to ribosomal proteins, various proteins from distinct components including intracellular organelles, cytosol, and cytoplasm were identified. Not surprisingly, membrane proteins were rarely reported, indicating that analyzing intact membrane proteins via TDP is still difficult when using standard conditions.

In biological process analyses (in Supplementary Table. 9), the most predominant GO term was *cytoplasmic translation*. Proteins involved in ribosomal subunit assembly were reported as abundant, and proteins associated with energy related GO terms were also found to be abundant (*ATP generation*, *ADP metabolic process*). In terms of molecular function (in Supplementary Table. 9), many proteins with binding functions including mRNA, rRNA, and ribosome binding were found. These results are highly consistent with the results from quantitative BUP analysis in Ishihama et al.^33^.

We then focused on the modifications in the proteoform IDs. TopPIC search allows for one modification per proteoform ID represented by a mass shift. Out of all proteoform IDs from FLASHIda runs, about 52% contained modifications. Fig. 3e and 3f show the frequent mass shifts found in the proteoform IDs (in both FI and ST datasets). Even if we did not specify any candidate modifications (or mass shifts) except for N-term modifications, most of the mass shifts could be explained by well-known modifications such as methylation, oxidation, and amino acid substitution. A large mass shift of +183 Da was also observed, which corresponds to artifacts from using AEBSF (Pefabloc) as a serine protease inhibitor during sample preparation^34^.

The remaining 48% of all proteoform IDs contained no mass shift, but many were truncated species. N-terminal methionine removal was present in about 30% of all proteoform IDs. In several proteins with well-documented signal peptides, we observed signal-peptide cleavage at the reported cleavage site (e.g., in metal-binding protein ZinT (UniProtKB: P76344) and inhibitor of g-type lysozyme (UniProtKB: P76002)). In the case of nickel/cobalt homeostasis protein RcnB (UniProtKB: P64534), the starting position of all proteoform IDs was one residue after the reported site (determined by Edman sequencing^35^), suggesting possible further processing after signal peptide cleavage. Supplementary Table 10 reports 419 proteoform IDs from top four selected proteins (two ribosomal proteins and two binding proteins; UniProtKB: P0ACF8, P0A7N1, P76344, P60438) with the four highest proteoform ID diversity. It also provides the manual validation results and possible interpretations for the modifications (see also Supplementary Fig. 20-24 for example annotations). In the two ribosomal proteins, common modifications were frequently observed such as oxidation and methylation. In addition, modifications like di-oxidation, di-methylation, and unexpected amino acid substitutions were also reported. In two binding proteins, many proteoform IDs were reported to have large mass shifts (ranging from 200 to 400 Da), representing bound metal ions or nucleotides as well as chemical adducts. To exclude chemical adducts and to further remove possible false modifications, we also ran TopPIC for FI90s and ST90s datasets with eight important candidate modifications including acetylation, phosphorylation, and formylation (see Methods and Supplementary File 1 for information on the eight candidate modifications). After the search, the proteoform IDs containing mass shifts not explained by the candidate modifications were discarded (see Supplementary Table 11 for the result). In this stringent search, the proteoform ID count in FI90s again outnumbered that in ST90s by 62% (976 vs. 602 on average). Moreover, the proteoform IDs with lysine acetylation were exclusively found in FI90s datasets.

## Discussion

The potential of IDA is beginning to emerge gradually, as seen in the development of many IDA methods for various MS-based applications such as TDP, BUP, and metabolomics^11, 25, 36–39^. The novel data acquisition algorithm presented here shows a drastic performance boost in terms of sensitivity and throughput for complex TDP samples. Since it does not require any change in the experimental set up, but is rather an extension module to the instrument software used for data acquisition, we believe that FLASHIda could be readily adopted for TDP studies to increase proteoform coverage at no additional cost.

Our algorithm currently focuses on the intelligent online selection of precursors, but it could obviously be improved to provide more fine control of the instrument, such as control of dissociation and ion injection duration tailored to the triggered precursor, in particular for high-mass proteoforms. While only HCD was used for this study, other dissociation methods, such as electron-transfer dissociation (ETD), could be dynamically selected for larger molecules to obtain better fragmentation of proteoforms^13^. The overhead to switch between dissociation methods could be compensated for by an increased identification rate of large proteoforms. Also the ion injection time could be adjusted based on the number of available precursor masses. From the data acquired for this study, we found that many of the MS spectra contain less than four candidate precursors owing to either sparse proteoform ions or low precursor qualities (see Fig. 3b and Supplementary Fig. 10). In such cases, increasing the injection time for MS2 generation or selecting multiple precursors for the same proteoform mass (of distinct changes) as in Lu et al.^11^ could lead to a higher number of proteoform IDs.

We also note that the precursor mass determination error and coeluted precursors would be one of the major issues in comprehensive TDP analysis and part of our future work. Many studies discussed FDR control in TDP (e.g., LeDuc et al.^40^), but this issue remains rarely discussed. We showed that such precursors lead to false proteoform discovery not measured nor controlled by conventional target-decoy approaches (Supplementary Fig. 2-3 and Supplementary Table. 1). While the use of precursor SNR is one of the effective solutions, other methods could be developed, e.g., using predicted RT of proteoforms, to minimize the precursor errors.

The current version of FLASHIda only supports proteoform identification by TopPIC due to the lack of interface between FLASHDeconv (for spectral deconvolution) and other proteoform identification tools like ProSight PC. Also, as stated above, the interface between FLASHIda and other deconvolution tools such as Xtract^41^ and TopFD could improve the identification performance. Thus, one of the future directions would be to develop such interfaces to easily deploy FLASHIda in various existing proteoform analysis pipelines.

While this study showcased the use of IDA for comprehensive TDP studies, different variations of the algorithm are currently under development for targeted proteoform analyses like global targeting^37^, deep characterization, and even *de novo* sequencing. For such applications, information on the target such as sequence, homologous proteoforms, composition, and candidate modifications could be used as inputs to FLASHIda for better acquisition tailored to the needs of a specific study. Furthermore, as FLASHIda enables the selection of interference-free precursors, it could be used to enhance the quantification accuracy of isobaric labeling-based proteoform quantification^42^. We anticipate that advanced data acquisition methods (to be) developed within FLASHIda would facilitate exploration of proteoform heterogeneity via TDP.

## Methods

### *E. coli* sample preparation

*E. coli* protein sample (Biorad, USA, cat#1632110) was dissolved in 0.1% formic acid solution to a final concentration of 1 g/L. Proteins were desalted with a solid-phase extraction tip packed with Poros R2 packing material (Thermo Scientific, USA) before the analysis.

### LC-MS/MS

A two-column setup was connected to EASY-nLC nano-HPLC (Thermo Scientific, San Jose, CA). Both the column (75 μm ID, 25cm) and the precolumn (100 μm ID, 3cm) were packed in-house using PLRP*-*S (1000Å, *5 μm)* packing material (Agilent, USA). Two different gradient lengths were used for both the standard and FLASHIda acquisition methods. Samples were recorded in triplicates for each condition, and 1 μg of the desalted sample was loaded for each run. Order of runs was randomized, and a column wash was included after each *E. coli* LC-MS/MS run to minimize the carryover. Mobile phase A consisted of 1% ACN 0.1% FA in water, and phase B of 95% ACN and 0.1% FA. In the 90 min runs, the concentration of buffer B was 5% for 5 min, 20% at 5 min, 55% at 70 min, 90% at 72 min, and 5% for the last 10 min. For the 30 min runs, it increased from 5% to 20% for the first two min, and was 55% at 19 min, 90% from 20 to 25 min, and 5% for the last 5 minutes.

Orbitrap Eclipse Tribrid mass spectrometer was used in the intact protein mode. Transfer tube temperature was set to 305 °C and 2.2 kV spray voltage was applied. Orbitrap resolution was set to 120,000 for MS1, and 60,000 for MS2 scans. Scan range was 500-2,000 and 400-2,000 Th for MS1 and MS2 scans respectively. HCD fragmentation was used with 29% normalized collision energy and 3 Th isolation window. Maximum injection time was 50 ms (MS1) and 500 ms (MS2) and AGC target was 200% (8e5) (MS1) and 1000% (5e5) (MS2). Only the precursors with charge states 4-50 were selected for fragmentation in both standard and FLASHIda runs. For standard runs, dynamic exclusion was applied with a duration of 30 sec and m/z tolerance of 1.5.

### FLASHIda software structure

FLASHIda is a console application written in C# (interaction with mass spectrometer) and C++ (deconvolution by FLASHDeconv, precursor selection, quality based exclusion, etc). It uses the instrument API (iAPI) provided by Thermo Scientific to communicate with the mass spectrometer. FLASHIda only can be used in iAPI compatible tribrid mass spectrometer series with instrument software version 3.4. The access to iAPI is available upon request from Thermo Scientific and is subject to a separate license. The parameters of acquisition (MS1 and MS2 scan parameters, and FLASHDeconv parameters) are controlled by a method file in XML format. Survey MS1 scans are received from the instrument through the iAPI in a vendor-specific format; briefly, the spectrum consists of an array of peak centroids and additional spectral metadata such as retention time, polarity, analyzer type, etc. Next, the spectra are converted into a format compatible with FLASHDeconv (separate m/z and intensity arrays) and sent to FLASHDeconv part via platform invocation (PInvoke). Upon receiving the spectrum, FLASHDeconv processes the spectrum and selects *N* (user controlled parameter) high quality precursors. The isolation window covers the complete precursor envelope range with 0.6 Th margin from both sides. The list of precursors (or isolation windows) returned by FLASHDeconv is used to create fragmentation scans in instrument specific format, which are added to a FIFO queue to be sent to the instrument. A survey MS1 scan as well as AGC ion trap scan is added to the queue after each set of fragmentation scans.

### QScore calculation via logistic regression

After the deconvolution, firstly per charge QScore, denoted as **Q**(*m, z*), is calculated for each precursor of (monoisotopic) mass *m* and charge *z* using a logistic regression. **Q**(*m*, *z*) is the probability that the MS2 spectrum from the charge *z* precursor of mass *m* is identified. The QScore of mass *m*, denoted as **Q**(*m*), is simply given by the maximum of **Q**(*m*, *z*) for all charges *z*. And the charge *z* such that **Q**(*m*, *z*)=**Q**(*m*) becomes the charge of the precursor.

Denote the mass of the proton by *c*, and the mass difference between ^13^C and ^12^C by δ. Given a mass *m* and charge *z*, if a peak has m/z of ((*m + n*δ)/*z*)+*c* for positive MS mode or m/z of ((*m + n*δ)/*z*)-*c* for negative mode within (user defined) tolerance, the peak is said to correspond to the *n*-th isotope (of mass *m* and charge *z*). Denote an isotope envelope of mass *m* and *z* as a vector **E**(*m*, *z*)=(*i_1_*, *i_2_*, …, *i_n_*), where *i_n_* denotes the summed intensity of the peaks corresponding to the *n*-th isotope. Suppose the charge range of this envelope is from *v* to *w*. Then the aggregated envelope **E**(*m*) is a vector given by adding all vectors **E**(*m*, *z*) for *z*=*v*,…,*w* element wise. **Q**(*m*, *z*) is furnished by a logistic regression with six features, two extracted from the charge *z* envelope, **E**(*m*, *z*), and four from the aggregated envelope **E**(*m*). The features are given by

1. Cosine similarity between the isotope pattern of **E**(*m*, *z*) and the theoretical one: charge_cos
2. Precursor SNR (signal to noise ratio) or SNR of **E**(*m*, *z*): precursor_SNR
3. Cosine similarity between the isotope pattern of **E**(*m*) and the theoretical one: mass_cos
4. Mass-level SNR or SNR of **E**(*m*): mass_SNR
5. Charge distribution score representing how evenly peak intensities are distributed along different charges: charge_dist
6. Average mass error of **E**(*m*) in PPM unit: mass_error

The theoretical isotope patterns are obtained by using averagine model^43^. The cosine similarity calculation is done as described in Jeong et al.^26^. The SNRs consider the peak locations as well as the shape of the isotope pattern for better separation of signal and noise components (see below). The charge distribution score calculation and average mass error calculation are described below as well. For cosine similarities and charge distribution scores *x*, we use *log*_2_(1+*x*) as feature values. For SNRs *y*, we use *log*_2_(1+*y*/(1+*y*)) as feature values. The mass error values are used as is. From the training of our logistic regression model, the weights for the QScore are determined by 0.4074, −1.5867, −22.1376, 0.4664, −0.4767, 0.541, and 20.248, for charge_cos, precursor_SNR, mass_cos, mass_SNR, charge_dist, mass_error, and intercept, respectively. From the weights, mass_cos is shown to be the most important feature for quality measure. Note that precursor intensity is not included in the feature set; our tests show that including it did not lead to any improvement in QScore (data not shown). Precursor mass is not used either since it may be strongly dependent on instrumental settings, e.g., resolution, making the model less robust than without it. To train the logistic regression model, six *E. coli* single run datasets were analyzed (deconvoluted by FLASHDeconv and identified by TopPIC at 1% spectrum and proteoform level FDR). The identified 13,205 precursors and unidentified 18,261 precursors were used as true and false classes, respectively. The training was done by Weka software (version 3.9.4)^44^ with default parameters for logistic regression. The input files to Weka software are found in Supplementary Table 12.

### SNR estimation

We first describe how the precursor SNR of the charge envelope **E**(*m*, *z*) is measured. Given a charge envelope **E**(*m*, *z*), denote the start m/z and the end m/z of **E**(*m*, *z*) as *t* and *s*, respectively. Out of all peaks between *t* and *s*, the peaks not corresponding to any isotope are noisy peaks and have only a noise component. The sum of squared intensities, or power, of these noisy peaks is written as *N_1_*. The peaks corresponding to isotopes mainly have a signal component of **E**(*m*, *z*), but also have a noise component from signal distortion, coelution, or simple thermal additive noise. This noise power (denoted by *N_2_*) and signal power (denoted by ***S***) within such peaks can be measured by solving the least square problem described below.

Denote the intensities of the theoretical isotopic distribution (from the averagine model^43^) of mass *m* by a vector **V**=(*v_1_*, *v_2_*, …, *v_n_*), where *v_i_* denotes *i-*th isotope intensity. Denote **E**(*m*, *z*) as **E** for simplicity. We want to know how much of the **V** component exists in **E**. To do so, we find a weight ***w*** such that ||**V**-***w*E**||^2^ is minimized. This is a typical least square problem and the solution for ***w*** is given by

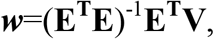

or equivalently,

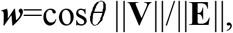

where cosθ is the cosine similarity between **E** and **V**, which is already available from FLASHDeconv. Once the weight ***w*** is calculated, the signal component power in **V** is given by ***S*** =||***w*E**||^2^=cos^2^θ ||**V**||^2^. The noise component power in **V** is given by ***N_2_*** =sin^2^θ ||**V**||^2^.

With the calculated signal power ***S*** and total noise power (***N_1_***+***N_2_***), the SNR is furnished by

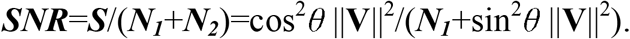

Note that this definition considers not only the noisy peaks outside the isotope locations but also those coexisting with the peaks corresponding to the isotopes.

The mass SNR for **E**(*m*) is done in the same way, except that the peaks of all charges are used in the place of charge *z* peaks for **E**(*m*, *z*). The calculation of SNR is implemented in FLASHDeconv without loss of computational efficiency.

### Charge distribution score calculation

The charge distribution score is calculated as follows. Given a mass *m*, let *i_z_* denote the summed intensity of the charge *z* peaks corresponding to any isotopes of the mass *m*. Further denote the minimum charge by *w* and the maximum charge *W*. Then for the mass *m*, we have a vector (*i_w_*, …, *i_W_*) that represents per charge intensity of *m*. Let *x* be the charge with the maximum per charge intensity, that is, the charge such that *i_x_* ≥ *i_z_* for all *z*=*w*,…,*W*. Then we define the left penalty *l_z_* for *z*=*w+*1,…,*x* by *l_z_* = max(0, *i_z_*_-1_ - *i_z_*) and the right penalty *r_z_* for *z*=*x*,…,*W*-1by *r_z_*=max(0, *i_z_*_+1_ - *i_z_*). The total penalty *P* is given by the summation of left and right penalties. Let *I* be the summation of all *i_z_* values. The charge distribution score is given by 1 - *P*/*I*.

### Average mass error calculation

Given the aggregated envelope **E**(*m*), the peaks corresponding to isotopes of **E**(*m*) may have different isotope indices. For a charge *z* peak of isotope index *n*, its monoisotopic mass *p* is calculated by *p=z*(*t* - *c*)-*n*δ, where *t* is the m/z of the peak, *c* denotes the mass of charge carrier (proton in positive MS and electron in negative MS), and δ the mass difference between ^13^C and ^12^C. Then its PPM error is given by 10^6^ |(*m* - *p*)/*m*|. The average mass error is given by the average value of these PPM errors over all the isotope corresponding peaks.

### TQScore (Total QScore) calculation

For a mass *m*, assume that it has been triggered *n* times with QScores *q_1_*,…,*q_n_*. TQScore of *m* is calculated by 1-(1 - *q_1_*)(1 - *q_2_*)…(1 - *q_n_*). This calculation assumes mutual independence among MS2 spectra of precursor mass *m* in terms of their identification rates. This is not rigorous in particular when multiple MS2 spectra are acquired from the precursors of the same charge state as they are expected to be highly similar to each other. However, since the primary goal of TQScore is not to estimate the identification rate accurately but to measure the (rough) quality of the mass for acquisition, we took this simple definition. To improve the estimation accuracy, methods taking the correlation between MS2 spectra into account should be applied, such as the one using Bayesian network.

### Target-decoy based FDR estimation of MS2 spectra of incorrect precursor masses

To test the effect of incorrect precursor masses on FDR estimation via target-decoy approaches, we took the deconvoluted spectra from FI90 dataset (by FLASHDeconv) and generated two sets of deconvoluted spectra with false precursor masses. The first set is generated by dividing all the precursor masses by two, simulating low harmonic artifacts (as they are the most common precursor errors from our observation). The second set was generated by adding arbitrary values ranging from −10 to −1 or from 1 to 10 to the precursor masses. Then these sets were identified using TopPIC at 1% spectrum and proteoform level FDR from the target-decoy approach. The results are shown in Supplementary Fig. 2-3 and Supplementary Table 1.

### FLASHDeconv commands for analysis of FI and ST datasets

#### For FI90 datasets

FLASHDeconv -in [input mzML] -out [feature deconvolution out tsv] -in_log [FLASHIda log file] -out_topFD [ms1 deconvoluted spectrum msalign] [ms2 deconvoluted spectrum msalign] -out_topFD_feature [ms1 feature] [ms2 feature] -min_precursor_snr 1.0 - Algorithm:min_charge 4 -Algorithm:max_charge 50 -Algorithm:rt_window 180 - Algorithm:min_mass 500 -Algorithm:max_mass 50000

#### For FI30 datasets

FLASHDeconv -in [input mzML] -out [feature deconvolution out tsv] -in_log [FLASHIda log file] -out_topFD [ms1 deconvoluted spectrum msalign] [ms2 deconvoluted spectrum msalign] -out_topFD_feature [ms1 feature] [ms2 feature] -min_precursor_snr 1.0 - Algorithm:min_charge 4 -Algorithm:max_charge 50 -Algorithm:rt_window 60 - Algorithm:min_mass 500 -Algorithm:max_mass 50000

#### For ST90 datasets

FLASHDeconv -in [input mzML] -out [feature deconvolution out tsv] -out_topFD [ms1 deconvoluted spectrum msalign] [ms2 deconvoluted spectrum msalign] -out_topFD_feature [ms1 feature] [ms2 feature] -min_precursor_snr 1.0 -Algorithm:min_charge 4 - Algorithm:max_charge 50 -Algorithm:rt_window 180 -Algorithm:min_mass 500 - Algorithm:max_mass 50000

#### For ST30 datasets

FLASHDeconv -in [input mzML] -out [feature deconvolution out tsv] -out_topFD [ms1 deconvoluted spectrum msalign] [ms2 deconvoluted spectrum msalign] -out_topFD_feature [ms1 feature] [ms2 feature] -min_precursor_snr 1.0 -Algorithm:min_charge 4 - Algorithm:max_charge 50 -Algorithm:rt_window 60 -Algorithm:min_mass 500 - Algorithm:max_mass 50000

The difference between FI and ST datasets is that in FI datasets, FLASHIda log file name is specified with the `in_log` option. The only difference between 90 and 30 min runs is the `rt_window` parameter value that decides the internal RT window size for FLASHDeconv. FLASHDeconv considers the spectra within this window for sensitive mass deconvolution (see Jeong et al.^26^). The generated msalign and feature files are used as the inputs to TopPIC.

### TopPIC commands for analysis of FI and ST datasets

#### For all datasets, we used the same command

`toppic.exe -d -t FDR -T FDR -u 16 [fasta] [ms2 msalign file name]`

For the search with candidate modifications (for Supplementary Table 11), we added the -i option as in: `toppic.exe -d -t FDR -T FDR -u 16 [fasta] [ms2 msalign file name] -i [modification text file name]`

The input modification text file for this search is given in Supplementary File 1.

Candidate modifications used in TopPIC search for analysis of FI90s and ST90s datasets to exclude chemical adducts and false positive modifications

| Name | Mass | Amino acid specificity | Position specificity | Unimod Acc. |
| --- | --- | --- | --- | --- |
| Acetylation | 42.010565 | K | any | 1 |
| Phosphorylation | 79.966331 | S,T,Y | any | 21 |
| Oxidation | 15.994915 | C,P,K,D,N,R,Y | any | 35 |
| Methylation | 14.01565 | C,K,R,H,D,E,N,Q | any | 34 |
| Di-oxidation | 31.989829 | M | any | 425 |
| Half of a disulfide bridge | -1.007825 | C | any | 374 |
| Formylation | 27.994915 | M | N-term | 122 |
| AEBS | 183.035399 | H,K,S | any | 276 |

Supplementary File1 is TopPIC input file for the above eight modifications.

## Data availability

The raw and converted mzML files for FI and ST triplicate datasets have been uploaded to MassIVE (https://massive.ucsd.edu) and are available under accession number MSV000087484 or under the digital object identifier https://doi.org/10.25345/C5FJ9G, where the identification results and MS1 signal screenshots (related to Supplementary Fig. 4-6, 16-18, and 21-24) are also found.

## Code availability

The C# source code of FLASHIda is published under a BSD three-clause license and is available at https://OpenMS.org/FLASHIda. The repository has detailed information on building and using FLASHIda. FLASHDeconv is implemented in C++ as a part of OpenMS^45^ and available as platform independent open-source software under a BSD three-clause license at https://OpenMS.org/FLASHDeconv.

## Acknowledgement

The authors acknowledge Jesse Canterbury and Shannon Eliuk from Thermo Fisher for their help with iAPI use. K.J. is thankful to Xiaowen Liu for insightful advice on the data analysis. Proteomics and mass spectrometry research at SDU are supported by generous grants to the VILLUM Center for Bioanalytical Sciences (VILLUM Foundation grant no. 7292 to O.N.J.) and PRO-MS: Danish National Mass Spectrometry Platform for Functional Proteomics (grant no. 5072-00007B to O.N.J). K.J., J.K., M.B., O.N.J., and O.K. acknowledge funding from the Horizon 2020 Marie Sklodowska-Curie Action ITN 2017 of the European Commission (grant 765502-A4B). K.J. and O.K. acknowledge EPIC-XS (project number 823839), funded by the Horizon 2020 programme of the European Union.

## Source data

Source Data for Fig. 2–3. FI and ST Proteoform IDs.

Source Data for Fig. 2–3. FI and ST Proteoform-spectrum matches (PrSMs).

## Supplementary information

Supplementary Figure 1. QScore distribution comparison between quality-based exclusion list and simple mass exclusion list in FI90 dataset.

Supplementary Figure 2. Numbers of identified spectra and proteoform IDs in ST90 dataset with intentionally introduced precursor mass errors.

Supplementary Figure 3. Analog of Fig. 3e for proteoform IDs of incorrect precursors.

Supplementary Figure 4-6. Selected precursors of low SNR (<1.0) from ST90 dataset.

Supplementary Figure 7. Analog of Supplementary Fig.2 for low precursor SNR identifications.

Supplementary Figure 8. Analog of Fig. 3e for low precursor SNR proteoform IDs.

Supplementary Figure 9. Screen capture of msconvert GUI displaying runtime parameters.

Supplementary Figure 10. The number of distinct precursor masses triggered in MS1 spectra.

Supplementary Figure 11. Scatter plots of identified and unidentified precursors on RT-mass and RT-m/z planes for FI90 and ST90 datasets.

Supplementary Figure 12. Mass distribution and identification rate for FI90 vs. ST90 (a) and FI30 vs. ST30 (b).

Supplementary Figure 13. Identification of FI and ST datasets with TopFD deconvolution.

Supplementary Figure 14. Analog of Supplementary Fig. 2 for low precursor SNR identifications with TopFD deconvolution.

Supplementary Figure 15. Analog of Fig. 3e for low precursor SNR proteoform IDs with TopFD deconvolution.

Supplementary Figure 16-18. Analog of Supplementary Fig. 4-6 for ST90 dataset with TopFD deconvolution.

Supplementary Figure 19. Proteoform level overlap coefficients among FI90s along summed intensity (a) and QScore (b).

Supplementary Figure 20. Distribution of the proteoform IDs (identified in FI90s datasets) of the selected four proteins (UniProtKB: P0ACF8, P0A7N1, P76344, P60438) in RT-mass plane.

Supplementary Figure 21. Analog of Supplementary Fig. 4 (a-f) and annotated sequence (g) for a DNA-binding protein H-NS (UniProtKB: P0ACF8) proteoform ID.

Supplementary Figure 22. Analog of Supplementary Fig. 4 (a-f) and annotated sequence (g) for a 50S ribosomal protein L31 type B (UniProtKB: P0A7N1) proteoform ID.

Supplementary Figure 23. Analog of Supplementary Fig. 4 (a-f) and annotated sequence (g) for a metal-binding protein ZinT (UniProtKB: P76344) proteoform ID.

Supplementary Figure 24. Analog of Supplementary Fig. 4 (a-f) and annotated sequence (g) for a 50S ribosomal protein L3 (UniProtKB: P60438) proteoform ID.

Supplementary Table 1. Identification results for ST90 dataset with intentionally introduced precursor mass errors.

Supplementary Table 2. Proteoform IDs of FI and ST datasets with TopFD deconvolution.

Supplementary Table 3. Proteoform IDs of FI and ST datasets with TopFD deconvolution and precursor SNR filtration.

Supplementary Table 4. Proteoform IDs of FI datasets with TopFD MS2 deconvolution (MS1 by FLASHIda).

Supplementary Table 5. PrSM identification results of FI and ST datasets with TopFD deconvolution.

Supplementary Table 6. PrSM identification results of FI and ST datasets with TopFD deconvolution and precursor SNR filtration.

Supplementary Table 7. PrSM identification results of FI datasets with TopFD MS2 deconvolution (MS1 by FLASHIda).

Supplementary Table 8. Proteins that are not identified in FI90s datasets out of previously reported 50 abundant E. coli proteins.

Supplementary Table 9. Gene ontology (GO) term analysis results for the identified proteins in FI datasets.

Supplementary Table 10. Proteoform IDs (identified in FI datasets) of the selected four proteins (UniProtKB: P0ACF8, P0A7N1, P76344, P60438) and possible interpretations thereof.

Supplementary Table 11. Proteoforms IDs (in FI90s and ST90s datasets) reported in TopPIC searches with eight candidate modifications.

Supplementary Table 12. Training dataset for QScore logistic regression.

Supplementary File 1. TopPIC input modification file for the eight modifications used in TopPIC searches for Supplementary Table 11.

