## Supplementary Note and Figures for "FLASHIda: Intelligent data acquisition for top-down proteomics that doubles proteoform level identification count"

**Supplementary Figure 1.**

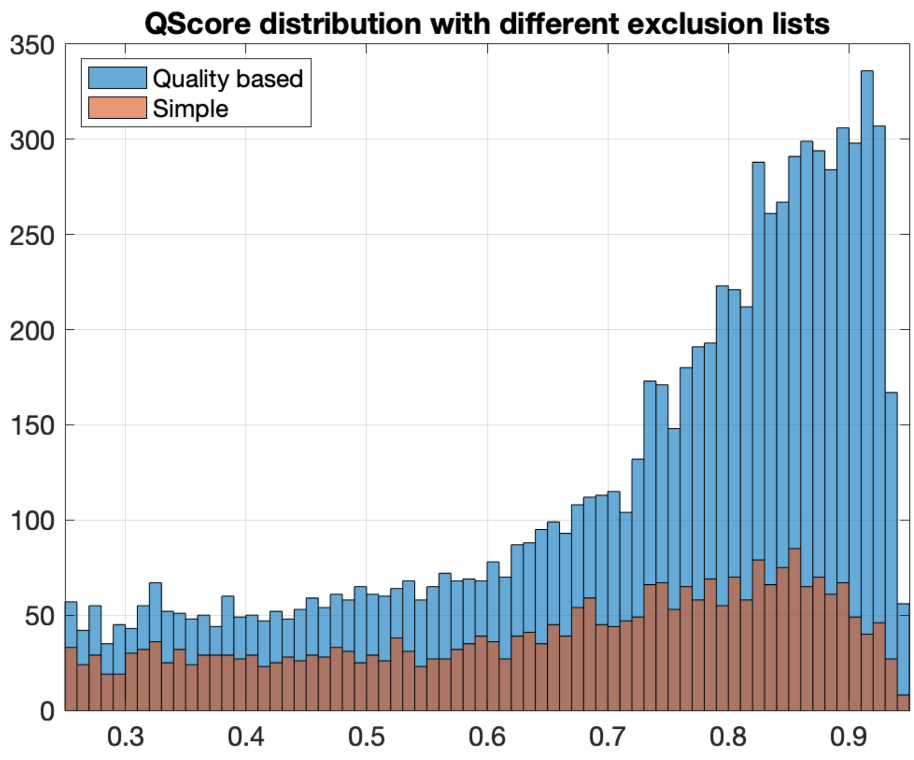


**Supplementary Figure 1. QScore distribution comparison between quality-based exclusion list and simple mass exclusion list in FI90 dataset.** Significantly more spectra have higher QScores with quality-based exclusion list than with simple mass exclusion list. The number of identified spectra can be estimated simply by adding up all QScores of spectra, because QScore is the estimated identification probability. The estimated number of identified spectra for quality-based exclusion is 6,096 while it is 1,884 for simple mass exclusion list. Note that the actual identified spectra for FI90 is 5,805.

**Supplementary Figure 2.**

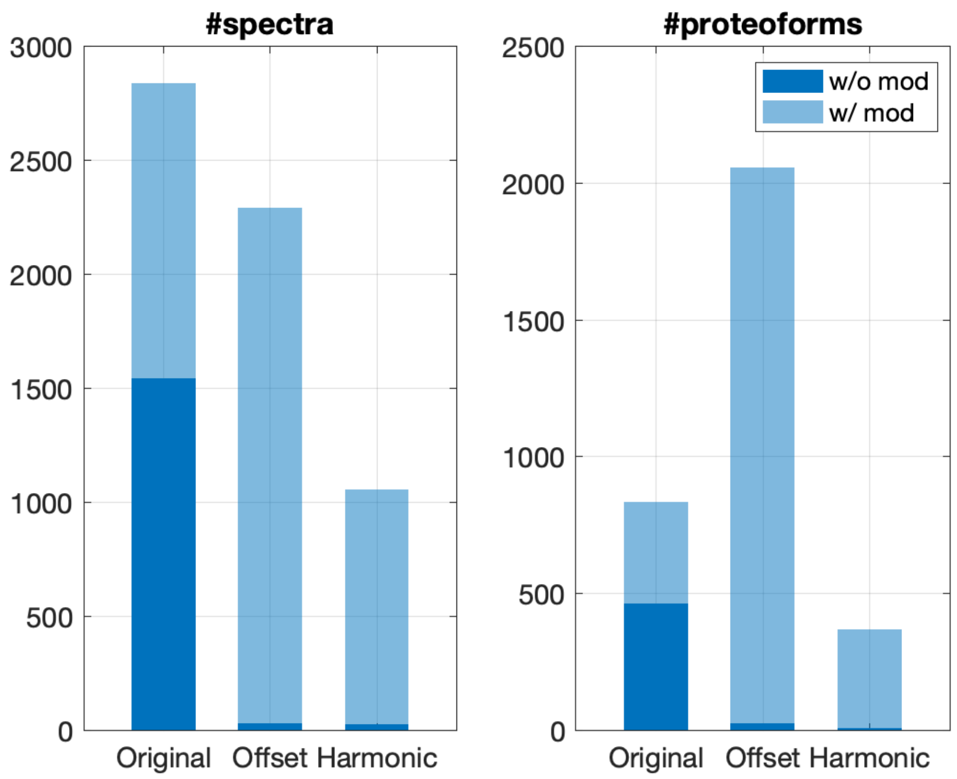


**Supplementary Figure 2. Numbers of identified spectra and proteoforms in ST90 dataset with intentionally introduced precursor mass errors.** The numbers of identified spectra and unique proteoforms from ST90 dataset are shown for three different cases: MS2 spectra of original precursor masses reported by FLASHDeconv (Original in x-axis label), original masses plus non-zero random offsets from 1 to 10 or -1 to -10 (Offset), and original masses divided by two simulating low harmonic artifacts (Harmonic). The bright blue portion indicates the portion of identification with an unknown modification. The identification was done with TopPIC at 1% spectrum and proteoform level FDR estimated by target-decoy database. While the original precursor masses resulted in the most identified spectra, the ones with offsets resulted in the largest number of proteoforms. This is because proteoforms with incorrect precursors usually are considered as “unique novel” proteoforms. In the harmonic simulation dataset as well, more than thousand spectra were identified (out of total 6,476), and hundreds of proteoforms were identified, even if most of them represent obvious false positives. Also, except for the original dataset, most identifications had unknown mass shifts (as expected from the spectra of incorrect precursor masses). This result clearly shows that incorrect precursor masses not only are hard to reduce via target-decoy approach but also may inflate the number of identified unique proteoforms. The identification results are found in Supplementary Table 1.

**Supplementary Figure 3.**

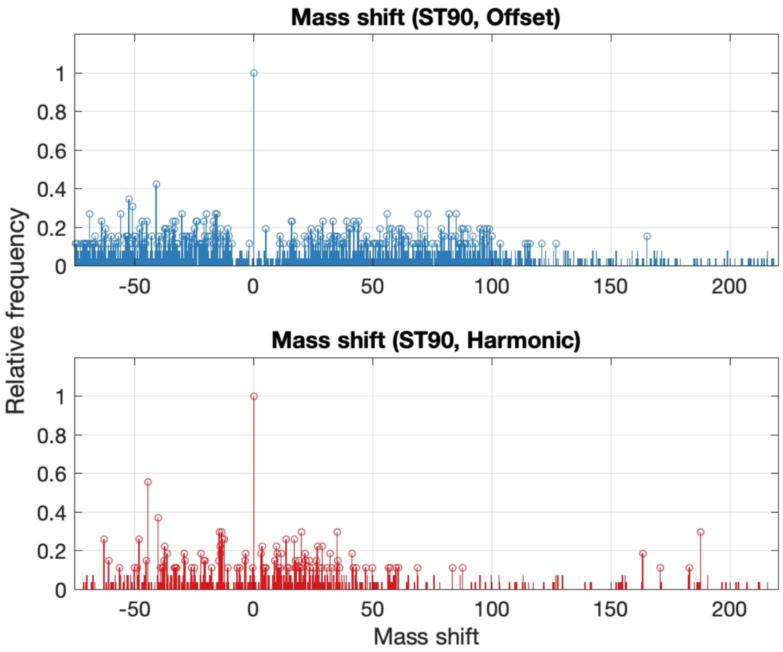


**Supplementary Figure 3. Analog of Fig. 3e for proteoform identifications of incorrect precursors.** The histograms of the mass shifts from the datasets in Supplementary Table 1 are shown. Top panel is for the precursors with random offsets (Offset in Supplementary Fig. 2) and bottom for the precursors divided by two (Harmonic in Supplementary Fig. 2). When compared to Fig. 3e, both histograms show significantly high levels of non-specific mass shifts, as expected from false positive hits.

**Supplementary Figure 4.**

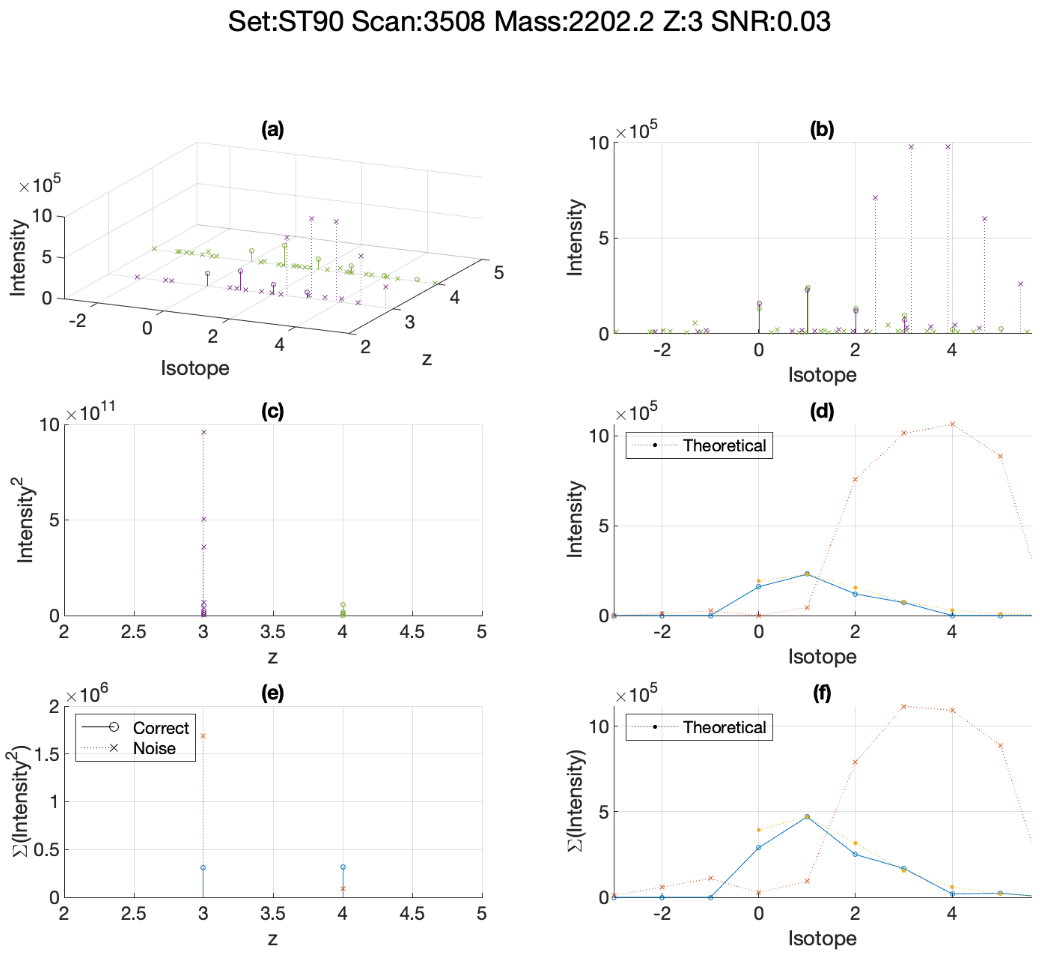


**Supplementary Figure 4. Selected precursors of low SNR (<1.0) from ST90 dataset.** The MS1 signals are shown for a representative precursor of low SNR. The figure title shows the dataset (Set:), scan number (Scan:), monoisotopic mass (Mass:), precursor charge (Z:), and precursor SNR (SNR:). **(a)** all peaks within isotopomer envelopes of distinct charges for the precursor monoisotopic mass. x-axis (Isotope) shows the isotope index, y-axis (z) shows the charge state, and z-axis (Intensity) the peak intensity. The circled peaks (or correct peaks) are the peaks corresponding to the isotopes of the mass, and crossed peaks (noise peaks) are the remaining peaks. **(b)** is the same as **(a)**, but viewed perpendicular to the x-y (isotope-intensity) plane, showing all peaks along isotope indices. **(c)** shows the squared intensity, or power, of individual peaks along charges. **(d)** shows the peak intensity along isotope indices only for the precursor charge (in this figure, z=3). All peaks are binned into the closest isotope indices. The blue (red) line shows the observed envelope from the correct (noise) peaks, and the yellow line is the theoretical envelope. **(e)** is drawn from **(c)** by aggregating the power of correct peaks (blue) and that of noise peaks (red) along charges. **(f)** is analog of **(d)** with the aggregated peak intensity for all charge states. The signal above is an example of coelution. **(a)** and **(b)** show explicit coelution (peaks for z = 3). In **(c)** and **(e)**, it is shown that noise power exceeds signal power for the precursor charge (z = 3). **(d)** and **(f)** show the presence of two distinct coeluted envelopes.

**Supplementary Figure 5.**

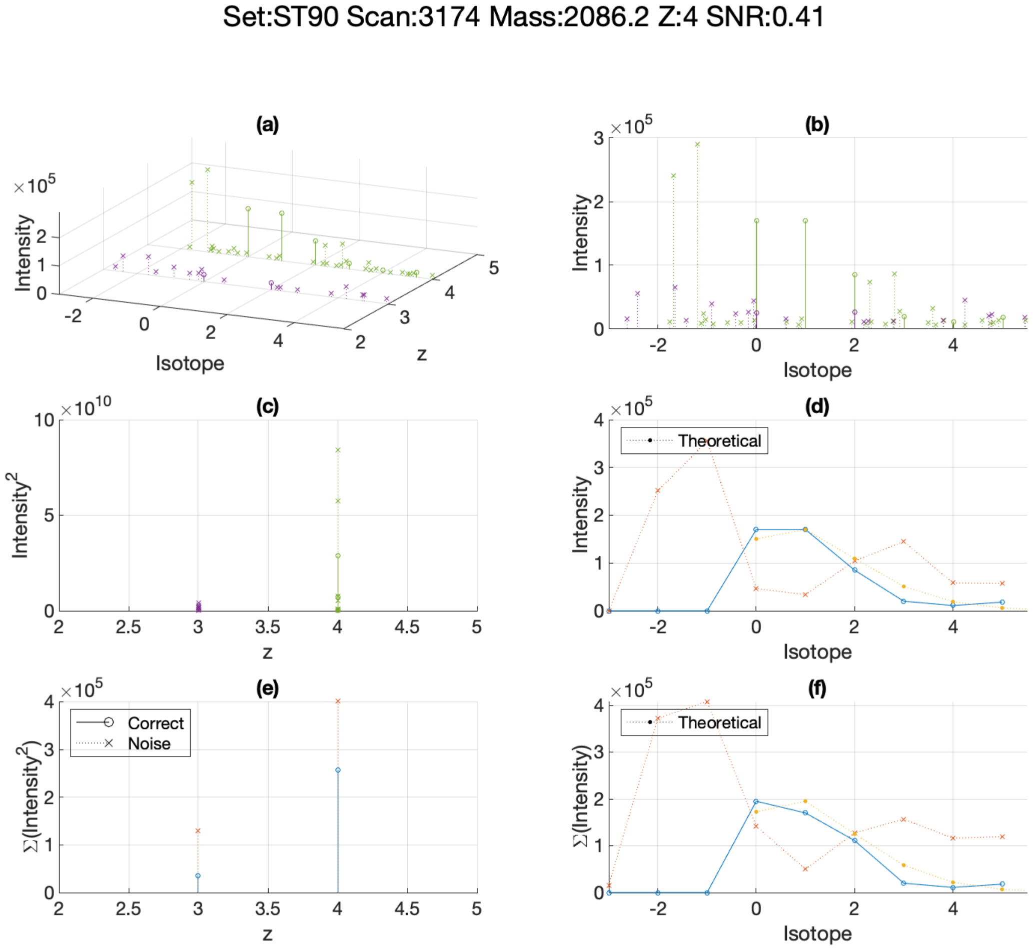


**Supplementary Figure 5. Selected precursors of low SNR (<1.0) from ST90 dataset.** Another example of coelution. Two envelopes of different charges are present within the range of the precursor envelope (z = 4).

**Supplementary Figure 6.**

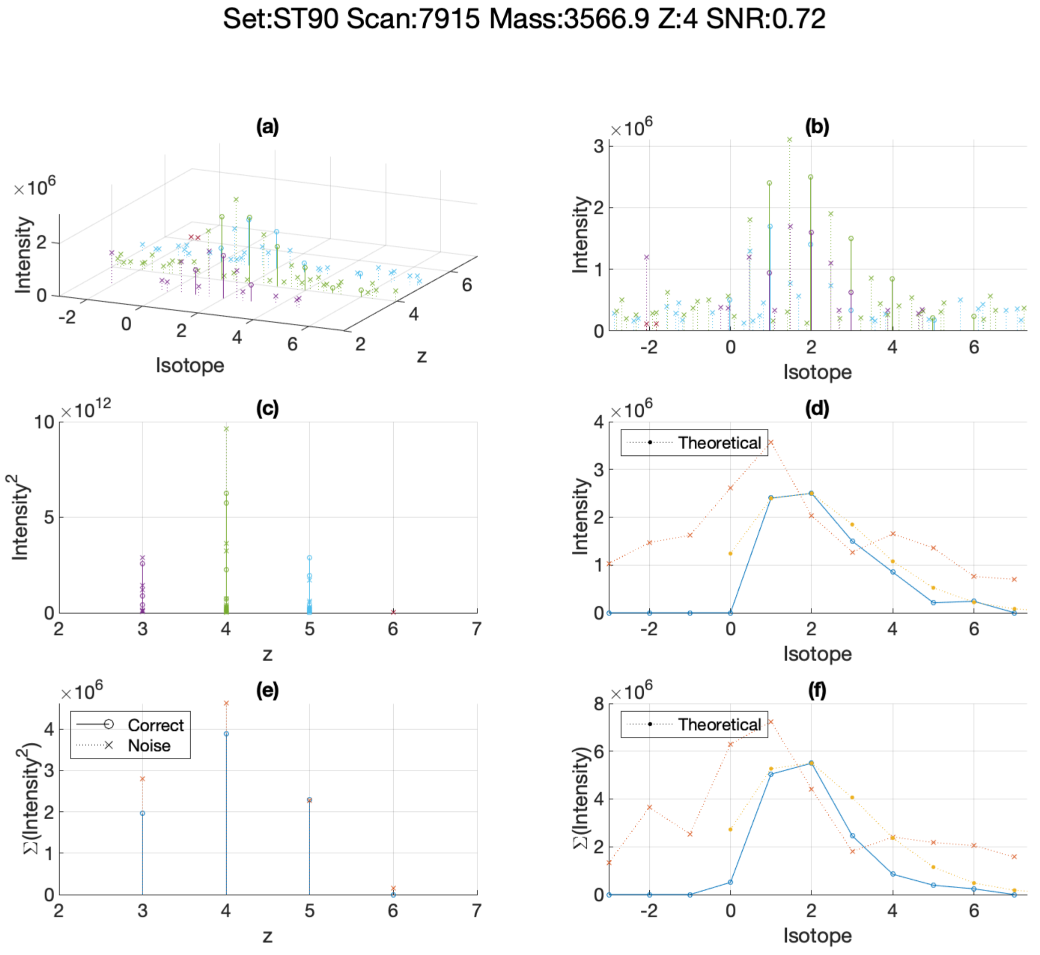


**Supplementary Figure 6. Selected precursors of low SNR (<1.0) from ST90 dataset.** This exemplifies the low harmonic mass artifact. The determined charge for the precursor is four, but from **(b)**, it is obvious that the correct precursor charge should be eight. The wrongful assignment of the precursor charge results in low SNR, which enables the removal of such artifacts. Signals of all low SNR precursors from ST90 dataset are deposited in MassIVE (https://massive.ucsd.edu) and are available under accession number MSV000087484 or under the digital object identifier https://doi.org/10.25345/C5FJ9G.

**Supplementary Figure 7.**

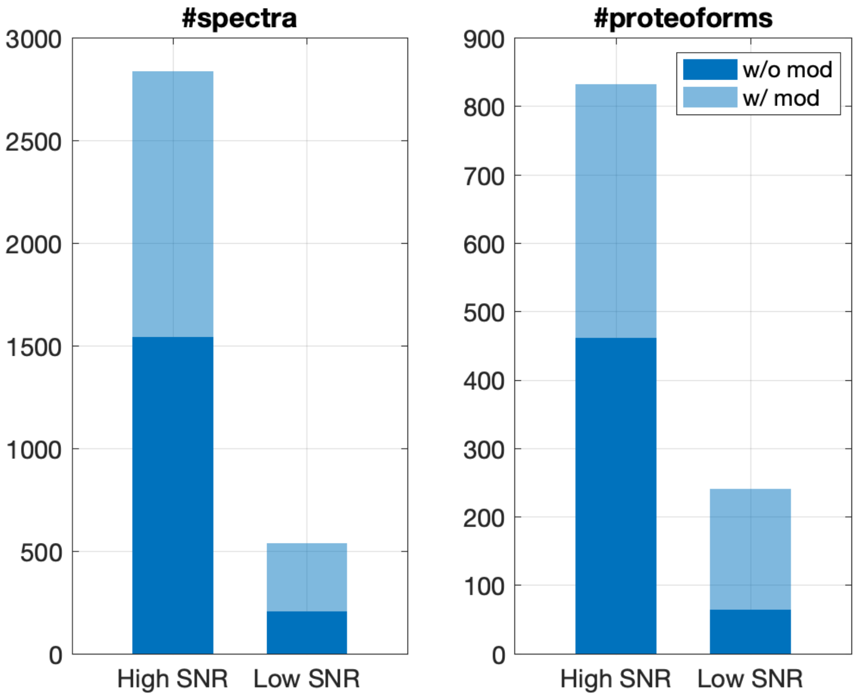


**Supplementary Figure 7. Analog of Supplementary Fig. 2 for low precursor SNR identifications.** ST90 dataset was identified without precursor SNR filtration, and the identified spectra and proteoforms were divided into two groups: from high-SNR precursors (precursor SNR>1; High SNR in x-axis) and from low-SNR precursors (precursor SNR<1; Low SNR). The left panel shows the number of identified spectra from each group and the right that of proteoforms. For the high SNR group, about 50% of spectral and proteoform identifications have no modification. But for the low SNR group, only 38% spectra and 27% proteoforms have no modification.

**Supplementary Figure 8.**

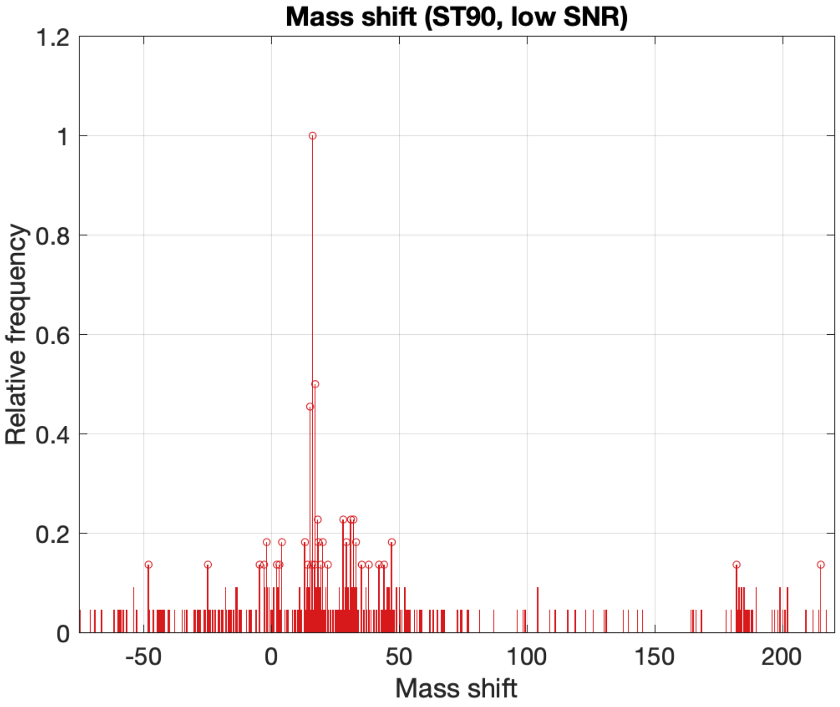


**Supplementary Figure 8. Analog of Fig. 3e for low precursor SNR proteoform identifications.** The histogram of the mass shifts of the proteoforms in the low SNR group in Supplementary Fig. 7 is shown. High level of non-specific mass shifts is observed when compared to the high SNR group in Fig. 3e bottom panel. Together with Supplementary Fig. 7, this result suggests that a large portion of proteoforms of low SNR precursors represent false positives.

**Supplementary Figure 9.**

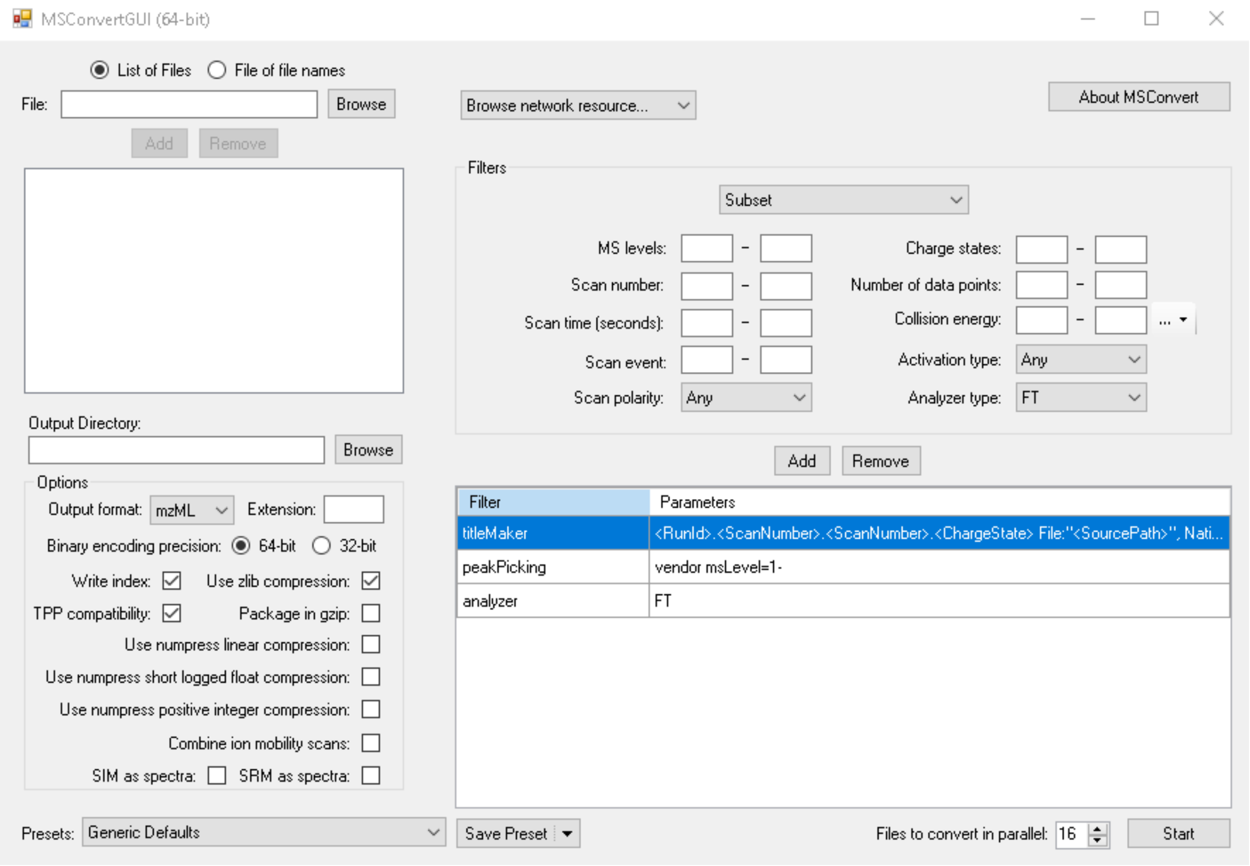


**Supplementary Figure 9**. **Screen capture of msconvert GUI displaying runtime parameters.** Thermo raw files containing profile mode spectra are converted into mzML format files by msconvert. All converted files are deposited in MassIVE (https://massive.ucsd.edu) and are available under accession number MSV000087484 or under the digital object identifier https://doi.org/10.25345/C5FJ9G.

**Supplementary Figure 10.**

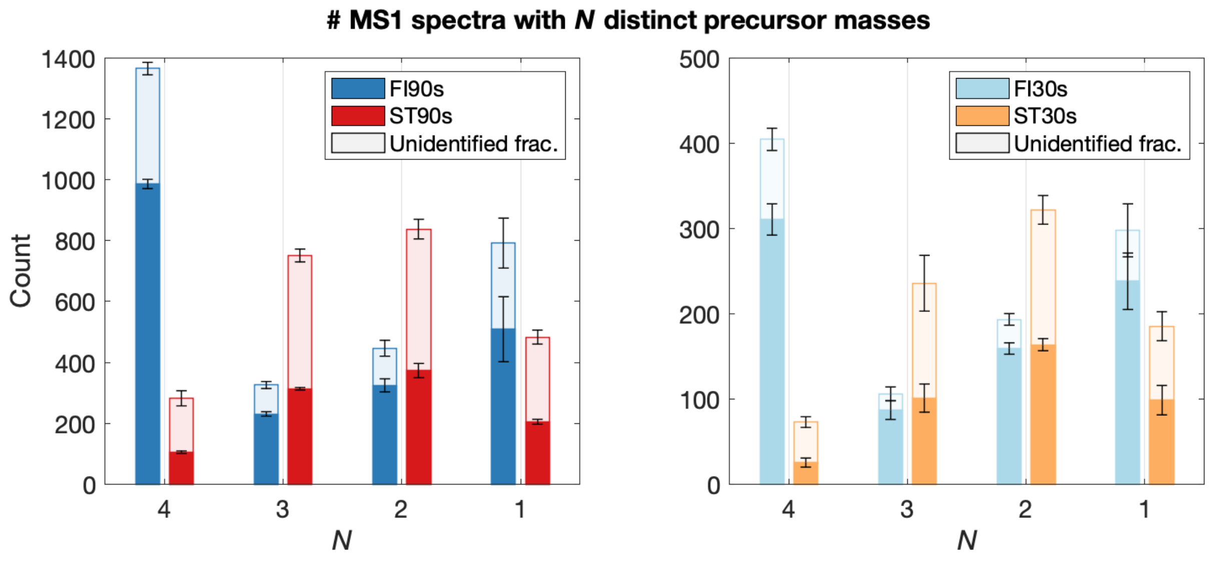


**Supplementary Figure 10. The number of distinct precursor masses triggered in MS1 spectra.** To more clearly observe if Top-4 QScore selection works as intended, we plot how many distinct masses were triggered per MS1 spectrum and how many out of them were identified. FLASHIda selected four distinct masses almost five times more often than the standard acquisition even though the numbers of total MS2 spectra from FLASHIda were about 20% less than from the standard for all datasets (seen in Fig. 2a, right panel). The portion of the identified spectra was also higher for FLASHIda than for the standard runs in all cases, regardless of the number of distinct masses

**Supplementary Figure 11.**

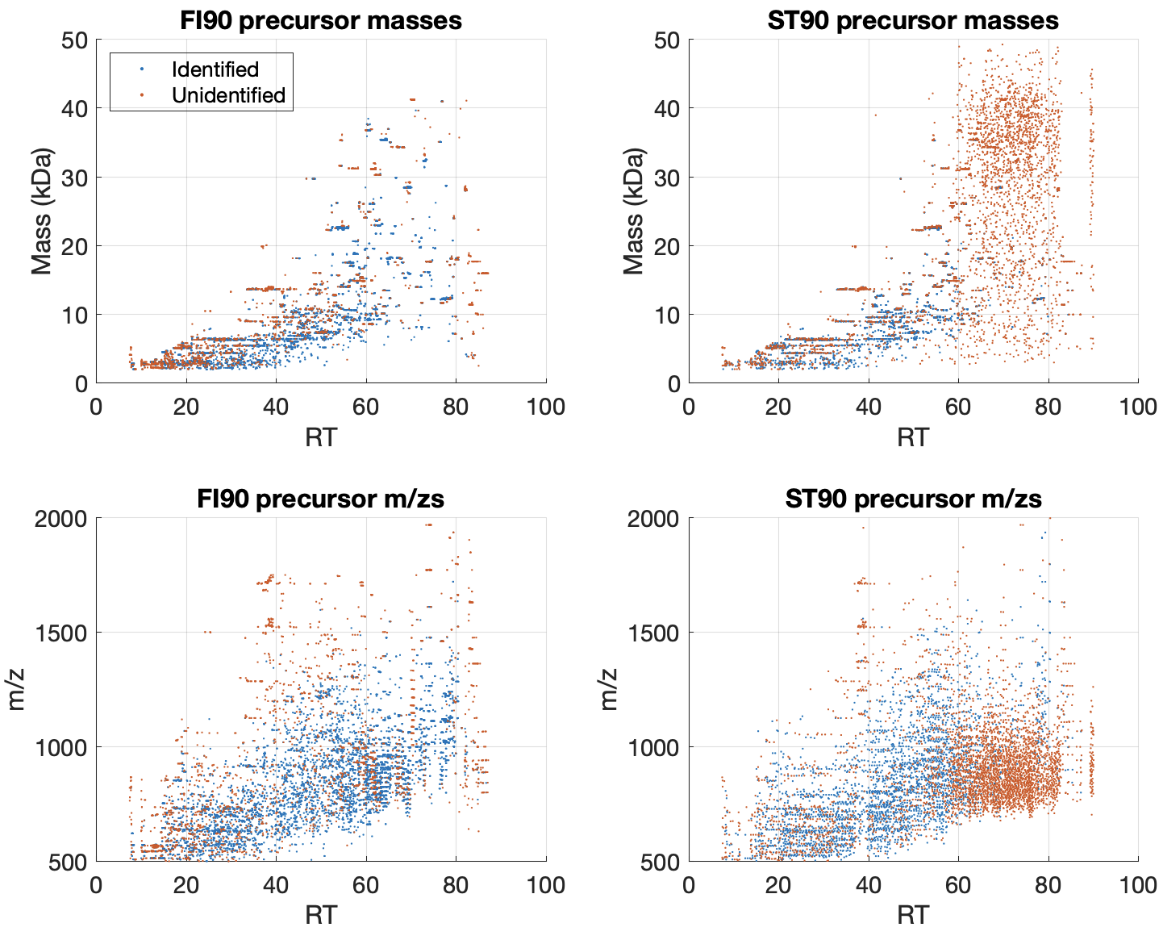


**Supplementary Figure 11. Scatter plots of identified and unidentified precursors on RT-mass and RT-m/z planes for FI90 and ST90 datasets.** The blue dots are the identified spectra and red unidentified. For both datasets, distributions on the RT-mass plane (upper panels) are far more sparse than on the RT-m/z plane (lower). In the 60-80 minute RT range, FLASHIda (left panels) did not acquire many MS2 spectra due to lack of high quality precursors. But the standard acquisition (right) generated many MS2 spectra in the same RT range, most of which resulted in unidentified spectra. While this shows the specific precursor selection of FLASHIda, it also shows that the identification of high mass proteoforms should be improved, for example, using other dissociations like ETD for high mass precursors.

**Supplementary Figure 12.**

**(a)**

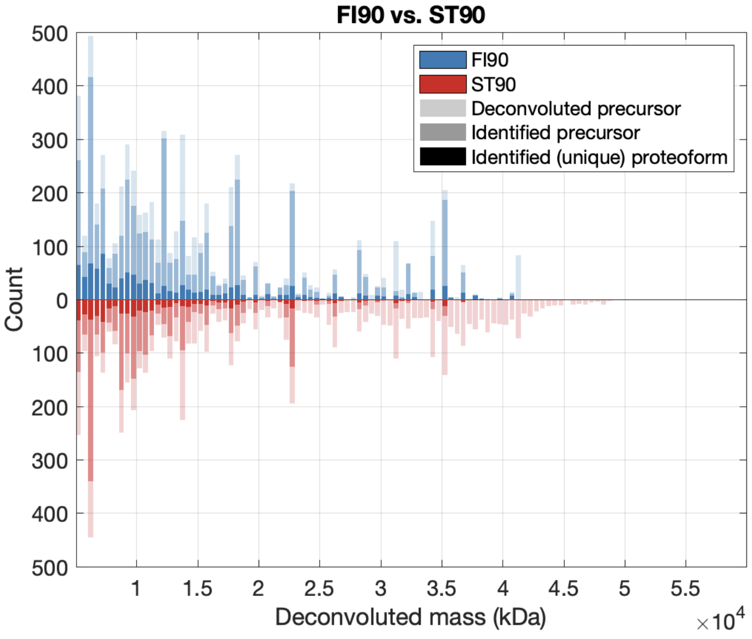


**(b)**

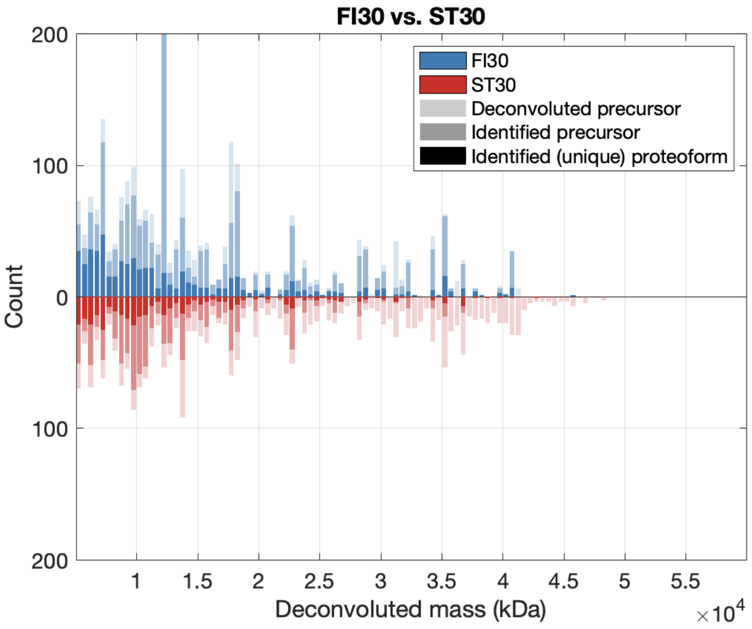


**Supplementary Figure 12. Mass distribution and identification rate for FI90 vs. ST90 (a) and FI30 vs. ST30 (b).** The blur bars are for FI and red for ST datasets. Each bar shows the number of deconvoluted spectra (high bright), identified spectra (medium bright), and identified unique proteoforms (dark). These plots show that even though the standard acquisition triggers more large masses (> 2 kDa) than FLASHIda does, FLASHIda identifies more high masses.

**Supplementary Figure 13.**

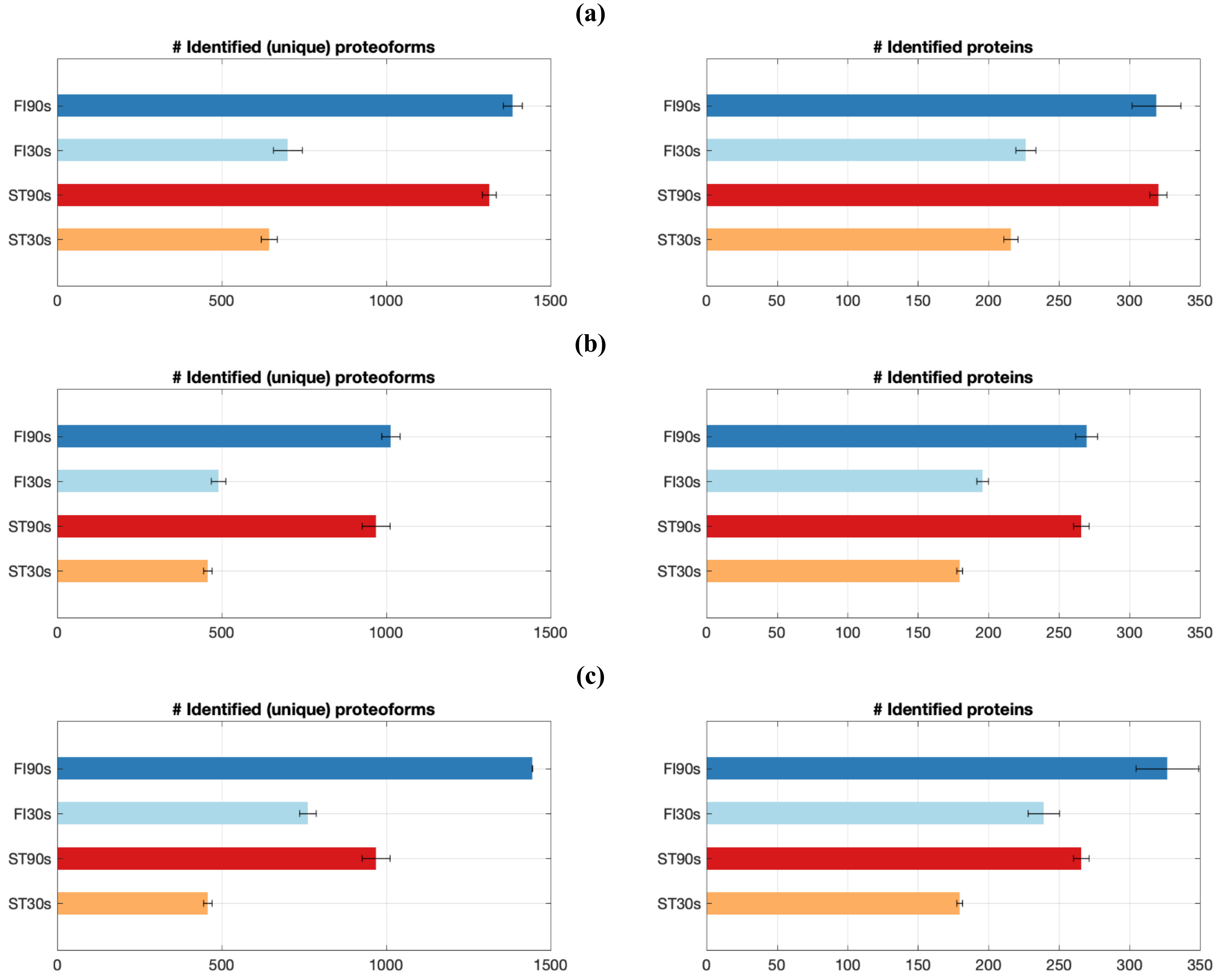


**Supplementary Figure 13. Identification of FI and ST datasets with TopFD deconvolution. (a)** the numbers of identified proteoforms (left) and proteins (right). FI and ST datasets were analyzed in the same way as Fig. 2a, except that the mass deconvolution (both MS1 and MS2) was done by TopFD instead of FLASHDeconv. **(b)** analog of **(a)** with precursor SNR filtration (SNR threshold = 1.0). The precursor SNRs were estimated using TopFD deconvoluted precursor masses. From the identified proteoforms, the ones with low precursor SNRs were discarded (see Supplementary Fig. 14-18 for the analysis of low SNR precursors). Note that an isolation window often contains multiple precursors of distinct precursor SNRs. Thus, even though FLASHIda selects an isolation window containing a precursor of high SNR, TopFD could select another precursor of low SNR from the same isolation window. Such precursors are filtered out for FIs datasets. **(c)** analog of **(b)** where only MS1 deconvolution was done by FLASHDeconv, or equivalently FLASHIda reported precursor masses were used. The boost from FLASHIda is not well observed when TopFD is used for MS1 deconvolution, regardless of the use of SNR filtering. But when FLASHIda reported precursor masses are used, the boost is again observed as shown in **(c)**.

**Supplementary Figure 14.**

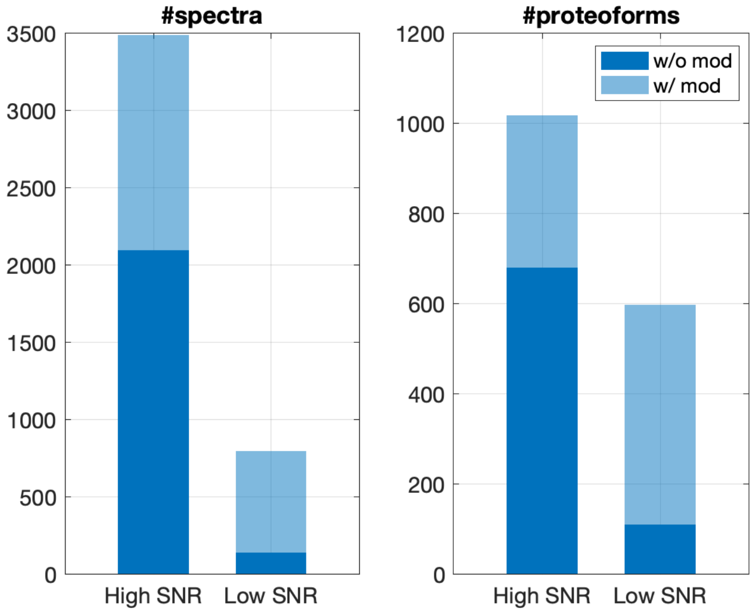


**Supplementary Figure 14. Analog of Supplementary Fig. 2 for low precursor SNR identifications with TopFD deconvolution.** We took the ST90 dataset analysis results from Supplementary Fig. 13 and divided them into two groups: ones of high-SNR precursors (precursor SNR>1; High SNR in x-axis) and the others of low-SNR precursors (precursor SNR<1; Low SNR). The left panel shows the number of identified spectra from each group and the right that of proteoforms. For the high SNR group, about 60% of spectral and 64% of proteoform identifications have no modification. But for the low SNR group, only 17% spectra and 18% proteoforms have no modification.

**Supplementary Figure 15.**

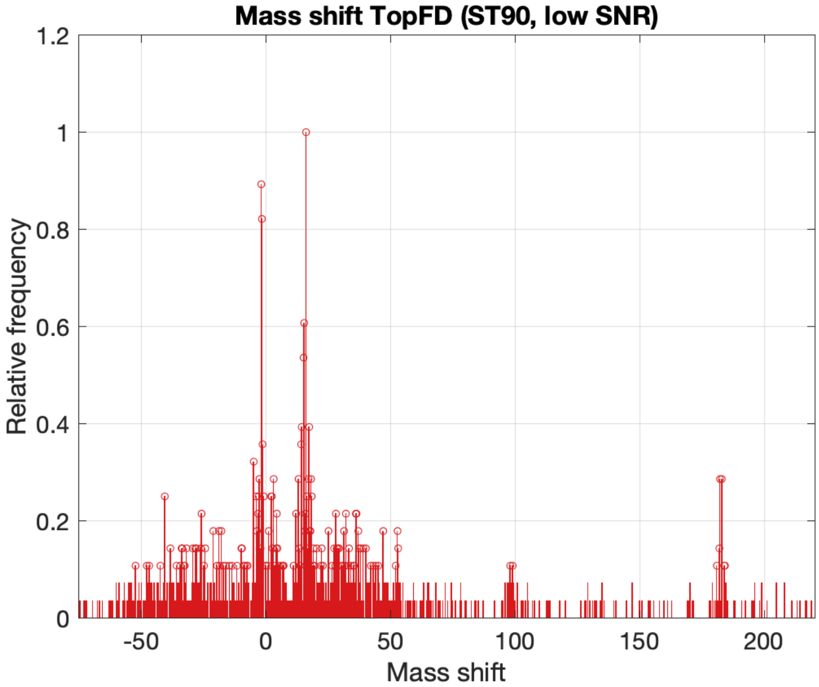


**Supplementary Figure 15. Analog of Fig. 3e for low precursor SNR proteoform identifications with TopFD deconvolution.** The histogram of the mass shifts from the low SNR group in Supplementary Fig. 14 is shown. As in Supplementary Fig. 8, a large number of non-specific mass shifts are observed. Together with Supplementary Fig. 14, this shows that a large portion of low precursor SNR proteoforms are false positives.

**Supplementary Figure 16.**

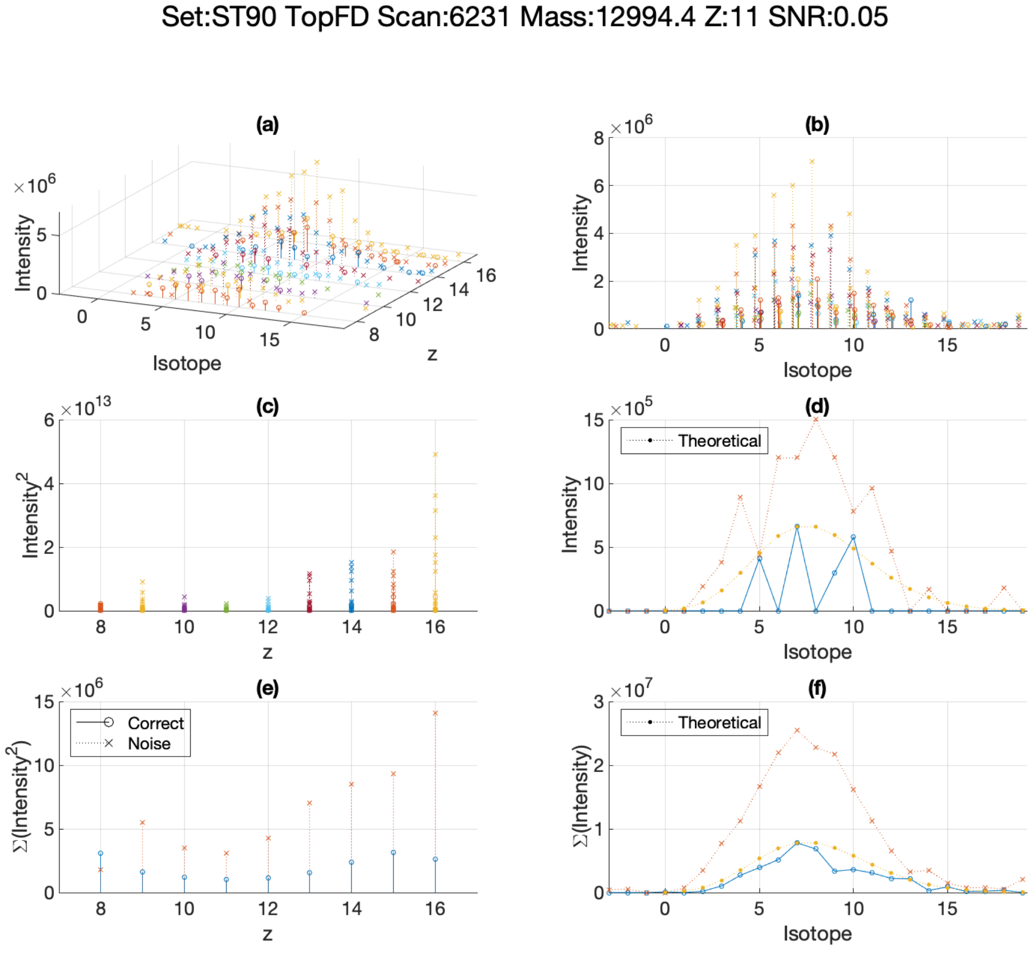


**Supplementary Figure 16. Analog of Supplementary Fig. 4 for ST90 dataset with TopFD deconvolution.** Signals of all low SNR precursors from ST90 dataset with TopFD deconvolution are deposited in MassIVE (https://massive.ucsd.edu) and are available under accession number MSV000087484 or under the digital object identifier https://doi.org/10.25345/C5FJ9G.

**Supplementary Figure 17.**

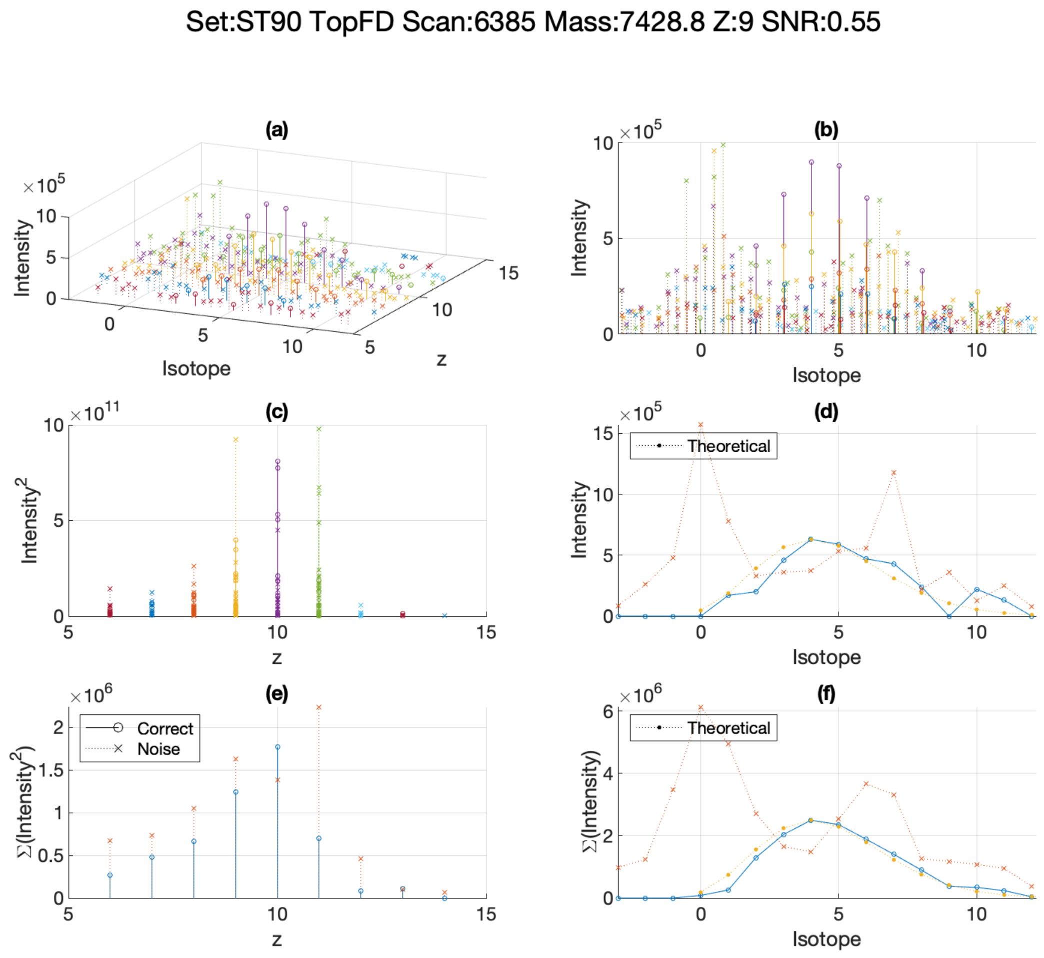


**Supplementary Figure 17. Analog of Supplementary Fig. 5 for ST90 dataset with TopFD deconvolution.**

**Supplementary Figure 18.**

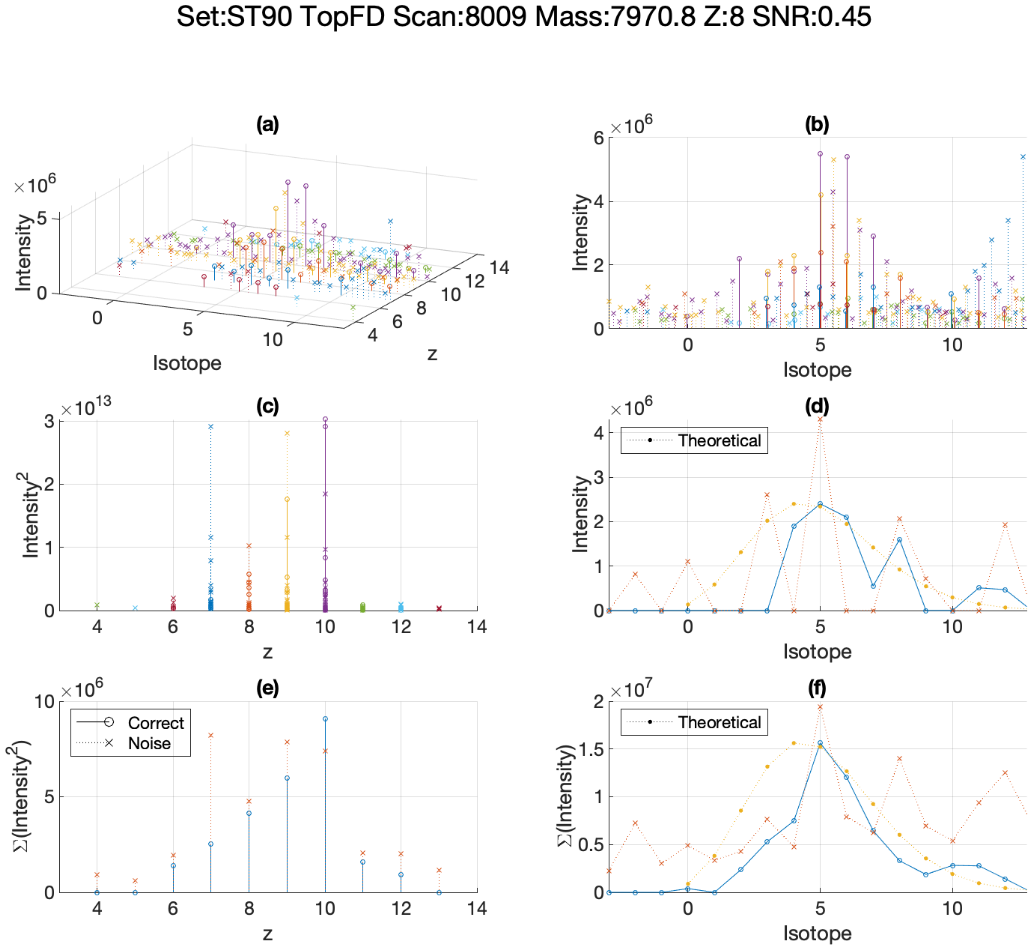


**Supplementary Figure 18. Analog of Supplementary Fig. 6 for ST90 dataset with TopFD deconvolution.**

**Supplementary Figure 19.**

**(a)**

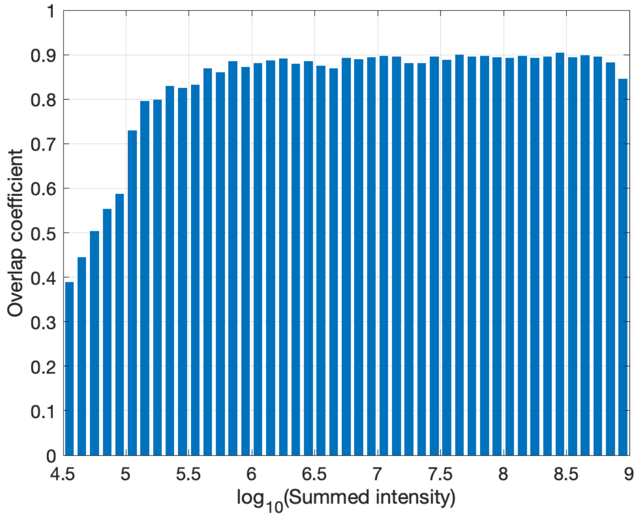


**(b)**

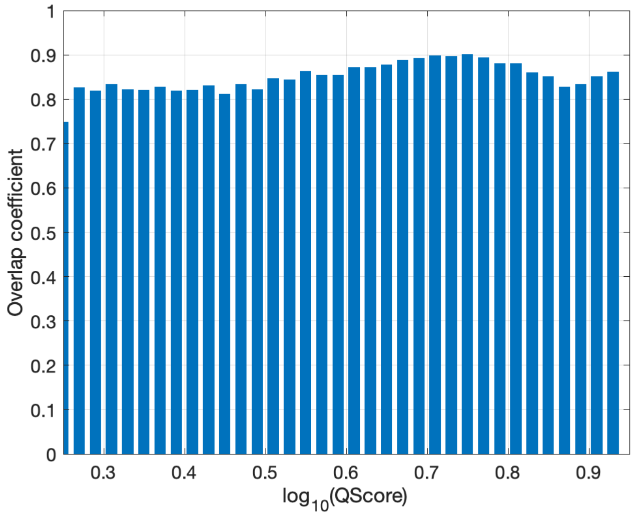


**Supplementary Figure 19. Proteoform level overlap coefficients among FI90s along summed intensity (a) and QScore (b).** Out of FI90s triplicates, possible three pairs were compared to each other (FI90 vs. FI90', FI90' vs. FI90", FI90" vs. FI90). Per each bin per pair, the overlap coefficient for the bin is calculated. Then the average overlap coefficient over three comparisons was drawn per bin. To calculate the overlap coefficient for a bin, the proteoforms corresponding to the bin from one set were collected. Then the collected ones were compared against all the proteoforms in the other set. In this way, the measurement is not affected by the QScore or intensity calculation consistency. But this calculation makes the overlap coefficients shown here (0.8-0.9) a bit higher than the overall overlap coefficient in Fig. 3a (0.63-0.65).

**Supplementary Figure 20.**

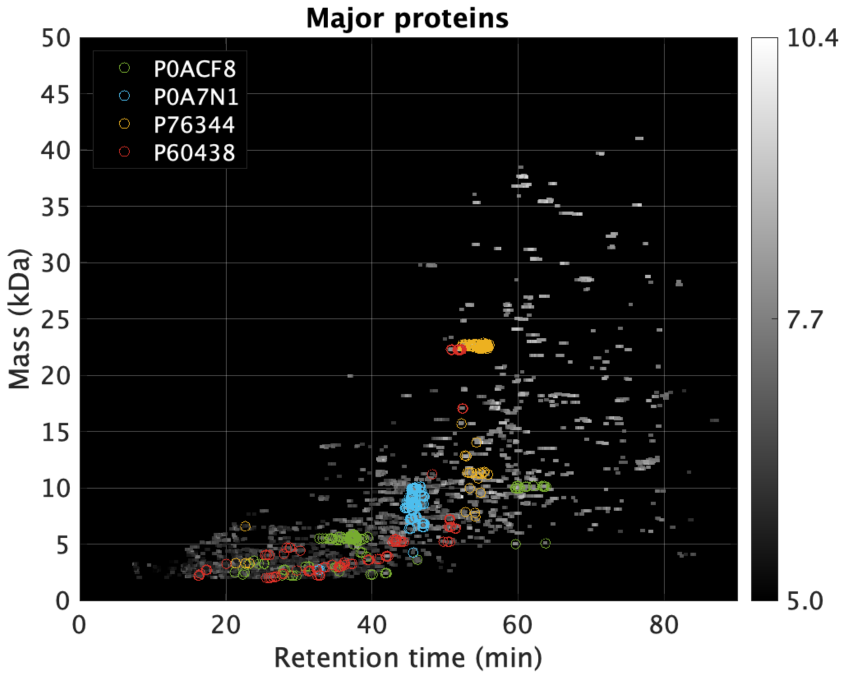


**Supplementary Figure 20. Distribution of the proteoforms (identified in FI90s datasets) of the selected four proteins (UniProtKB: P0ACF8, P0A7N1, P76344, P60438) in RT-mass plane.** Four proteins with highest proteoform heterogeneity have been selected from FI90s datasets: DNA-binding protein H-NS (UniProtKB: P0ACF8), 50S ribosomal protein L31 type B (UniProtKB: P0A7N1), metal-binding protein ZinT (UniProtKB: P76344), and 50S ribosomal protein L3 (UniProtKB: P60438). The proteoforms are color coded on the RT-mass plane along with other proteoforms in gray.

**Supplementary Figure 21.**

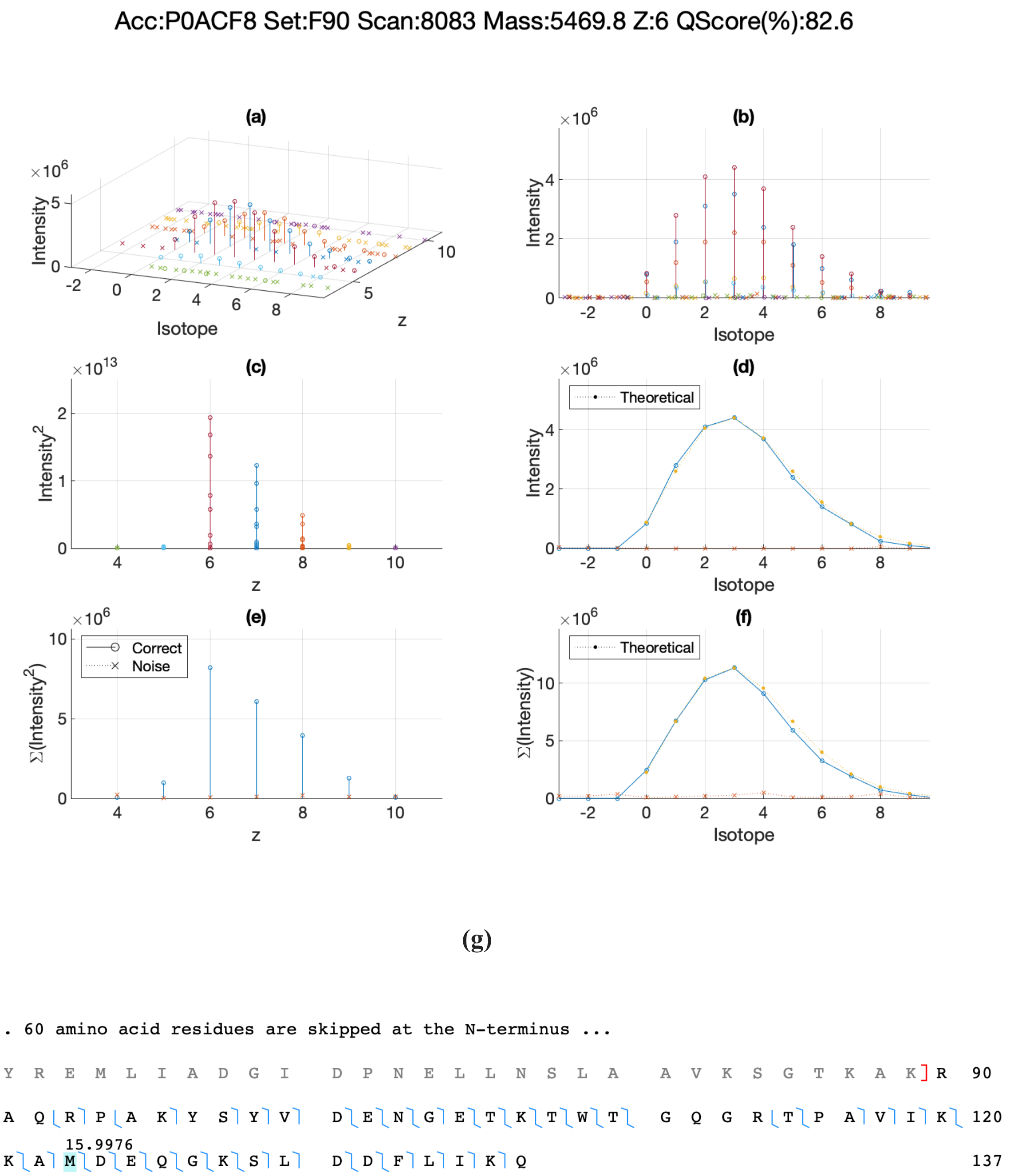


**Supplementary Figure 21. Analog of Supplementary Fig. 4 (a-f) and annotated sequence (g) for a DNA-binding protein H-NS (UniProtKB: P0ACF8) proteoform.** MS1 signals of all proteoforms from the major four proteins are deposited in MassIVE (https://massive.ucsd.edu) and are available under accession number MSV000087484 or under the digital object identifier https://doi.org/10.25345/C5FJ9G.

**Supplementary Figure 22.**

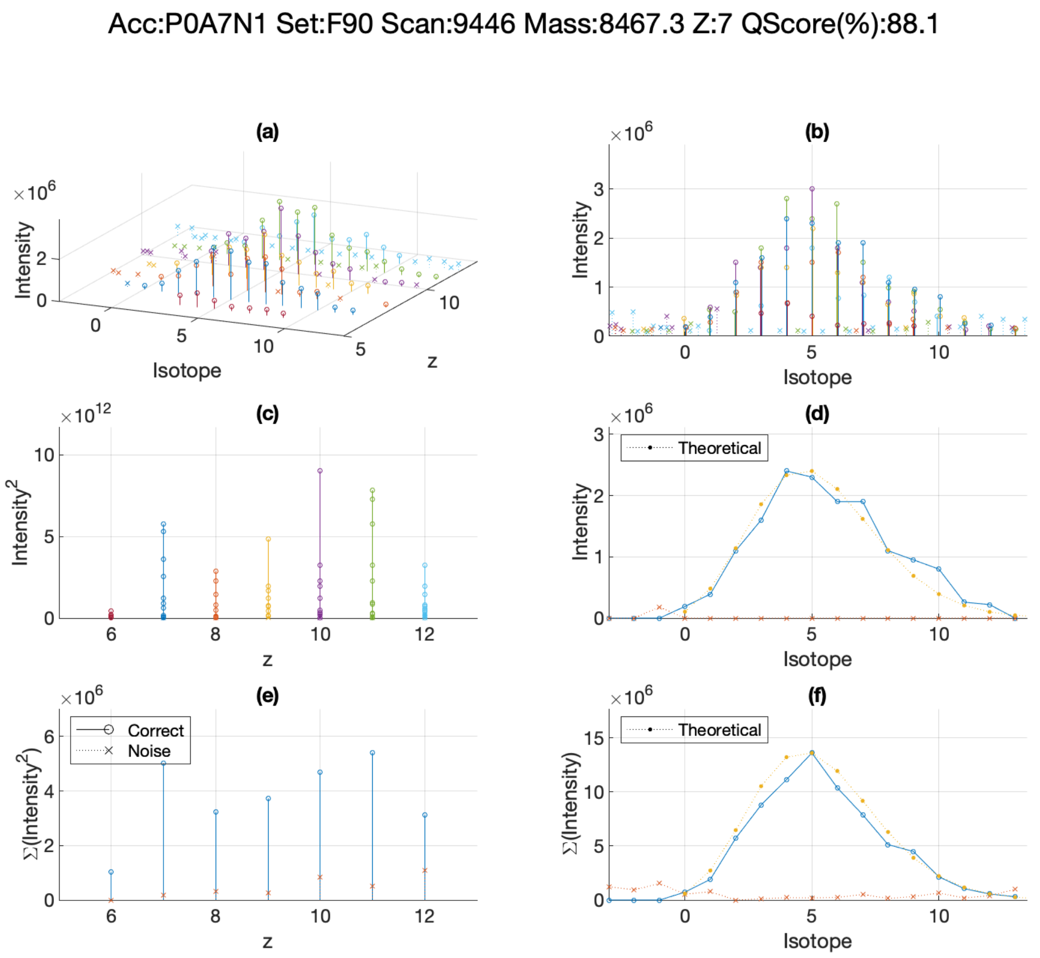


**(g)**

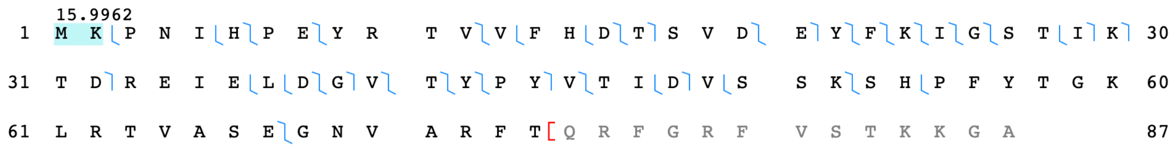


**Supplementary Figure 22. Analog of Supplementary Fig. 4 (a-f) and annotated sequence (g) for a 50S ribosomal protein L31 type B (UniProtKB: P0A7N1) proteoform.**

**Supplementary Figure 23.**

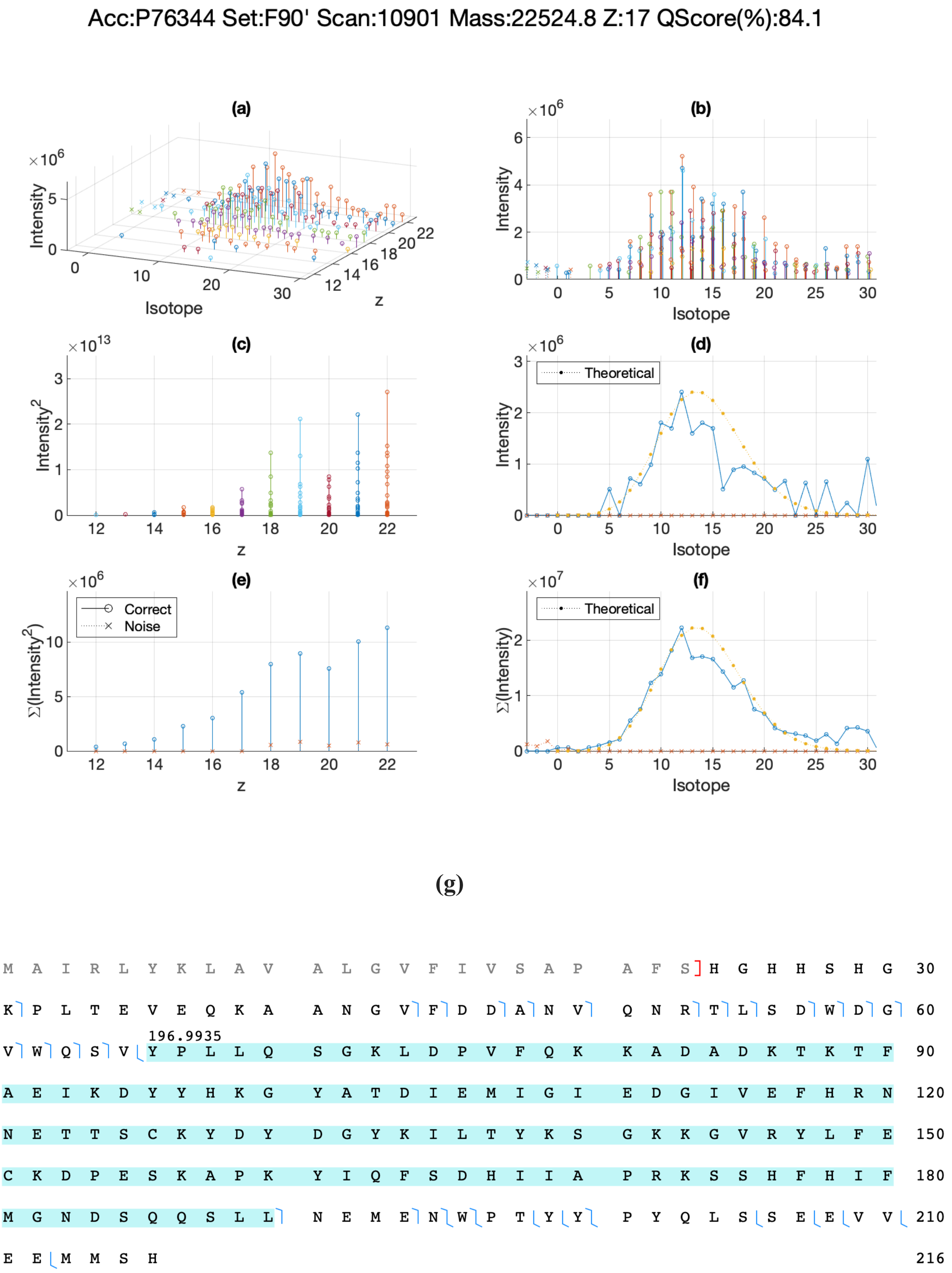


**Supplementary Figure 23. Analog of Supplementary Fig. 4 (a-f) and annotated sequence (g) for a metal-binding protein ZinT (UniProtKB: P76344) proteoform.**

**Supplementary Figure 24.**

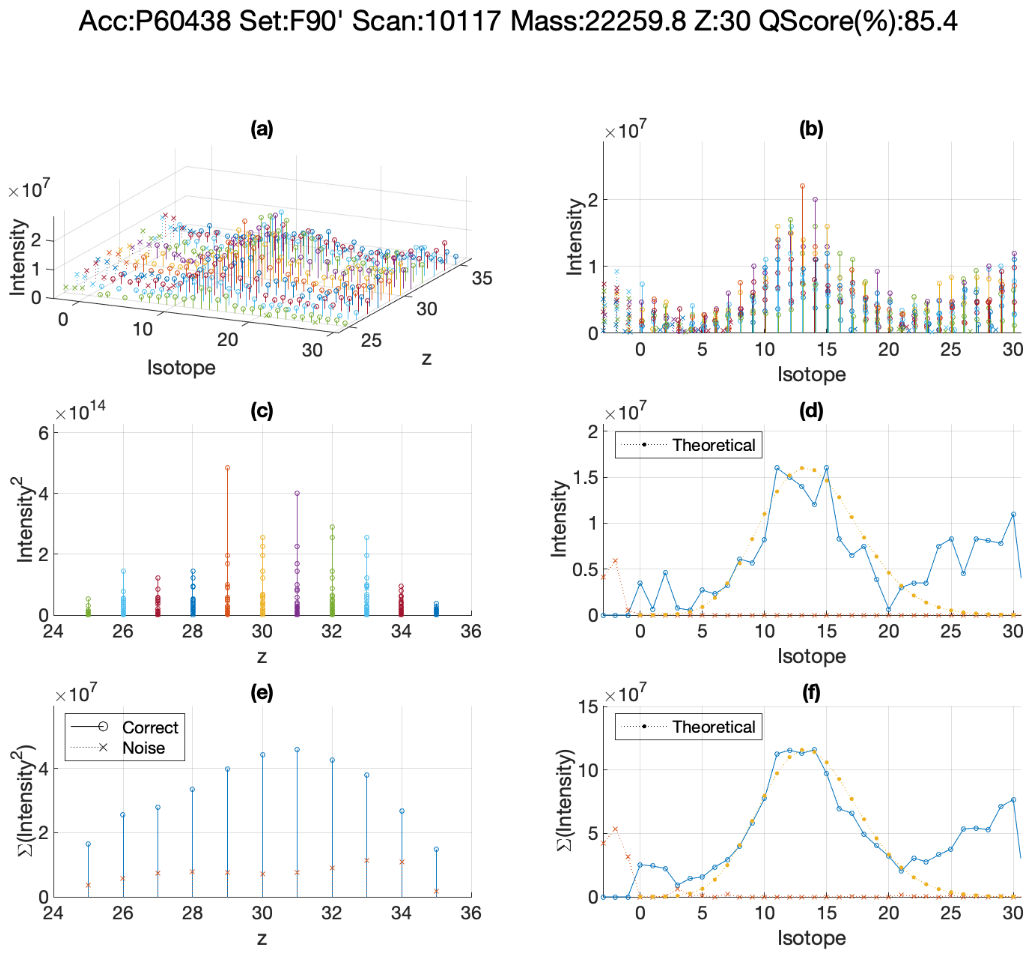


(g)

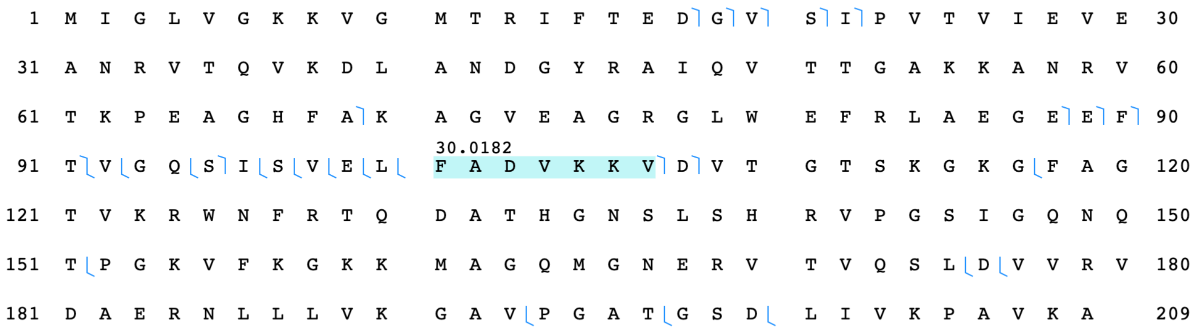


**Supplementary Figure 24. Analog of Supplementary Fig. 4 (a-f) and annotated sequence (g) for a 50S ribosomal protein L3 (UniProtKB: P60438) proteoform.**

**Supplementary Table 1. Identification results for ST90 dataset with intentionally introduced precursor mass errors.**

**Supplementary Table 2. Proteoform identification results of FI and ST datasets with TopFD deconvolution.**

**Supplementary Table 3. Proteoform identification results of FI and ST datasets with TopFD deconvolution and precursor SNR filtration.**

**Supplementary Table 4. Proteoform identification results of FI datasets with TopFD MS2 deconvolution (MS1 by FLASHIda).**

**Supplementary Table 5. PrSM identification results of FI and ST datasets with TopFD deconvolution.**

**Supplementary Table 6. PrSM identification results of FI and ST datasets with TopFD deconvolution and precursor SNR filtration.**

**Supplementary Table 7. PrSM identification results of FI datasets with TopFD MS2 deconvolution (MS1 by FLASHIda).**

**Supplementary Table 8. Proteins that are not identified in FI90s datasets out of previously reported 50 abundant *E. coli* proteins.**

**Supplementary Table 9. Gene ontology (GO) term analysis results for the identified proteins in FI datasets.**

**Supplementary Table 10. Proteoforms (identified in FI datasets) of the selected four proteins (UniProtKB: P0ACF8, P0A7N1, P76344, P60438) and possible interpretations thereof.**

**Supplementary Table 11. Proteoforms identified (in FI90s and ST90s datasets) in TopPIC searches with eight candidate modifications.**

**Supplementary Table 12. Training dataset for QScore logistic regression.**

**Supplementary File 1. TopPIC input modification file for the eight modifications used in TopPIC searches for Supplementary Table 11.**
